# The influence of autotoxicity on the dynamics of vegetation spots

**DOI:** 10.1101/2020.07.29.226522

**Authors:** Annalisa Iuorio, Frits Veerman

## Abstract

Plant autotoxicity has proved to play an essential role in the behaviour of local vegetation. We analyse a reaction-diffusion-ODE model describing the interactions between vegetation, water, and autotoxicity. The presence of autotoxicity is seen to induce movement and deformation of spot patterns in some parameter regimes, a phenomenon which does not occur in classical biomass-water models. We aim to analytically quantify this novel feature, by studying travelling wave solutions in one spatial dimension. We use geometric singular perturbation theory to prove the existence of symmetric, stationary and non-symmetric, travelling pulse solutions, by constructing appropriate homoclinic orbits in the associated 5-dimensional dynamical system. In the singularly perturbed context, we perform an extensive scaling analysis of the dynamical system, identifying multiple asymptotic scaling regimes where (travelling) pulses may or may not be constructed. We show that, while the analytically constructed stationary pulse corresponds to its numerical counterpart, there is a discrepancy between the numerically observed travelling pulse and its analytical counterpart. Our findings indicate how the inclusion of an additional ODE may significantly influence the properties of classical biomass-water models of Klausmeier/Gray–Scott type.

## 1. Introduction

Self-organized vegetation patterning has been proved in recent years to play an important role in our understanding of climate change and catastrophic shifts [44], in particular in arid and semi-arid environments (e.g., [9, 10]). Several reasons have been hypothesised to explain the occurrence of such patterns, focusing on spatial interactions between vegetation and the abiotic environment (e.g., [11, 20, 23, 27, 45, 49, 50]). The main mechanisms that have been identified in vegetation self-organization rely on the transport of water and nutrients toward vegetated patches through a scale-dependent (short-range positive, long-range negative) feedback, allowing plants to act as ecosystems engineers [16, 32]. Water availability alone, however, fails to explain the emergence of vegetation patterns in environments where water availability is not limited (see [46] and references therein).

A mechanism that is by now well established in the description of vegetation dynamics is plant-soil negative feedback (see, e.g., [15, 34, 42]). This feedback, which was already well-known in agriculture since ancient time, underlies the practice of “crop rotation” after repeated monoculture to avoid “soil sickness” - a situation where the same plant species cannot be grown within the same region after a certain amount of time. The origin of this feed-back has been linked to several biological phenomena, such as the presence of soilborne pathogens, the changing composition of soil microbial communities [31], and the accumulation of autotoxic compounds from decomposing plant litter [4, 39]. In particular, it has been proved that plant-soil negative feedback occurs because of inhibitory effects of extracellular DNA [40]. Extensive studies have revealed that this negative feedback plays an important role in several aspects of vegetation dynamics, such as species coexistence and biodiversity [38], as well as spatial organisation of plants by means of clonal rings [5, 6] and patterns [35].

In [35], in particular, the authors have found that the presence of autotoxicity induces – for some parameter ranges, and in agreement with experimental observations – the occurrence of dynamic patterns, with a non-symmetric distribution of biomass. The governing process is represented by the attempt of the biomass to “escape” areas with high levels of toxicity. The analytical foundation for such ecologically relevant results, however, is still lacking.

In this paper, we aim to take a first step in analytically justifying the numerical findings in [35] by using recently developed analytical techniques [14, 48]. In particular, we prove existence of both stationary and travelling pulses in one spatial dimension. To this aim, we use a convenient scaling, which allows us to transform the model introduced in [35] into an extension of the classical Klausmeier/Gray-Scott (KGS) model. For this classical KGS model, a large number of extensive analytical and numerical studies have been performed, investigating the existence, stability and dynamics of several types of patterns [12, 13, 28, 29, 41, 49, 50]; note that these references are but an incomplete selection of the vast literature on pattern formation in the KGS model.

We show that the extension of the KGS model with a third, spatially homogeneous, ordinary differential equation that models autotoxicity, can significantly influence the development and behaviour of pattern solutions, inducing the formation of travelling pulses which have been proven not to exist in the corresponding toxicity-free case [13].

From an analytical point of view, the inclusion of an additional model component significantly increases the complexity of the model, and hence introduces a potential obstacle for the successful analysis of pattern formation in this extended KGS model. One of the main results of this paper is that several asymptotic scaling regimes can be identified where the techniques developed in a two-component setting [14] nevertheless can be extended to analyse the existence of pulse patterns in this new three-component ‘reaction-diffusion-ODE’ model.

In the original KGS model, the existence and stability of pulse solutions sensitively depends on the asymptotic scaling of model components and parameters, see e.g. [2]. In order to analytically understand certain numerically observed behaviour, the ‘proper’ asymptotic scaling regime has to be identified. In the classical KGS model, this has already proven to be a subtle problem (see e.g. [48]); the inclusion of an additional model component exacerbates this subtility by increasing the number of scaling regimes that can be considered. Since choosing a particular asymptotic scaling is, to a certain extent, a prerequisite for the application of the analytical techniques presented in [14], the in-depth investigation of asymptotic scaling regimes relevant for investigating the existence of stationary and travelling pulse solutions, is a major part of this paper.

The paper is structured as follows: in Section 2, we present the original (dimensional) biomass-water-autotoxicity model [35], and the nondimensionalised, scaled model version we use in our subsequent analysis. Section 3 revolves around a collection of numerical results obtained by simulating both the original model in two spatial dimensions, and the rescaled, extended KGS model in a one-dimensional domain, showing the influence of autotoxicity. Section 4 is devoted to the existence for stationary and travelling pulses. Finally, a discussion of the presented results and an outline for future work in Section 5 conclude the paper.

## 2. The model

The original model introduced in [35] describes the dynamics of biomass (*B*), water (*W*), and autotoxicity (*T*) on a two-dimensional spatial domain, all measured in kg m^−2^, as follows: Growth of biomass *B* is mediated by water availability, its intrinsic mortality, and the toxic compounds; availability of water *W* is affected by precipitation, evaporation, and transpiration (plant water uptake); finally, autotoxicity *T* grows due to the decomposition of dead plants and is removed from the soil by intrinsic degradation and precipitation. In mathematical terms, this corresponds to

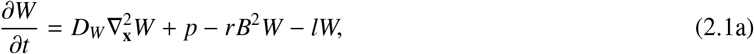

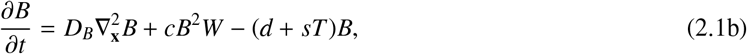

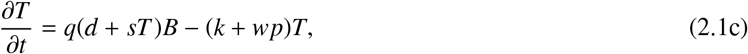

with time *t* ≥0 measured in days (d) and space **x** = (*x, y*) ∈ ⊂ ∇ Ω ℝ^2^ in meters (m); 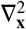 is the usual planar Laplace operator. System (2.1) is an extension of the classical Klausmeier model [27] on flat terrain, hence the absence of an advection term in (2.1a) (see e.g. [26]). A detailed description of the model together with a biological interpretation of System (2.1) is given in [35]. For an overview of the model parameters as in [35] see Table 1: these values are either chosen in accordance with [27] and [6] or selected from within an order-of-magnitude feasibility range.

**Table 1:** Model parameters for model (2.1) as presented in [35]. Note (*): the dimension of *w* was stated incorrectly in [35].

| Parameter | Description | Units | Value in [35] |
| --- | --- | --- | --- |
| $c$ | Growth rate of $B$ due to water uptake | $\text{m}^4\text{d}^{-1}\text{kg}^{-2}$ | $2 \cdot 10^{-3}$ |
| $d$ | Death rate of biomass $B$ | $\text{d}^{-1}$ | $1 \cdot 10^{-2}$ |
| $k$ | Decay rate of toxicity $T$ | $\text{d}^{-1}$ | $1 \cdot 10^{-2}$ or $2 \cdot 10^{-1}$ |
| $l$ | Water loss due to evaporation or drainage | $\text{d}^{-1}$ | $1 \cdot 10^{-2}$ |
| $p$ | Precipitation rate | $\text{kg d}^{-1}\text{m}^{-2}$ | $[0, 2]$ |
| $q$ | Proportion of toxins in dead biomass | – | $5 \cdot 10^{-2}$ |
| $r$ | Rate of water uptake | $\text{m}^4\text{d}^{-1}\text{kg}^{-2}$ | $3.5 \cdot 10^{-1}$ |
| $s$ | Sensitivity of plants to toxicity $T$ | $\text{m}^2\text{d}^{-1}\text{kg}^{-1}$ | $0$ or $2 \cdot 10^{-1}$ |
| $w$ | Washing out of toxins by precipitation | $\text{m}^2\text{kg}^{-1}(\ast)$ | $1 \cdot 10^{-3}$ |
| $D_B$ | Diffusion coefficient of biomass $B$ | $\text{m}^2\text{d}^{-1}$ | $1 \cdot 10^{-2}$ |
| $D_W$ | Diffusion coefficient of water $W$ | $\text{m}^2\text{d}^{-1}$ | $8 \cdot 10^{-1}$ |

We rescale and nondimensionalise (2.1) as follows. First, we define the nondimensional space and time variables

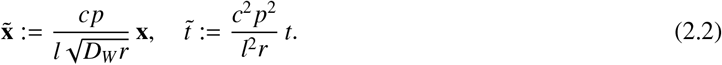

Next, we define the nondimensional model components

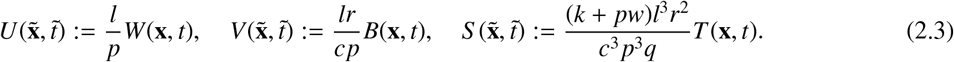

The model in the new components (*U, V, S*) reads

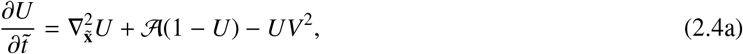

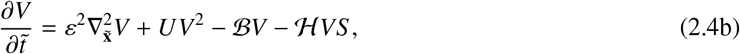

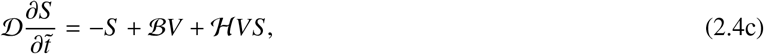

defined on a two-dimensional spatial domain 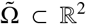 for 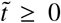 0. The (nondimensional) parameters in (2.4) are defined as

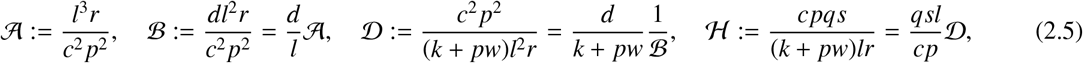

while the (square root of the) diffusivity ratio gives rise to the natural small parameter

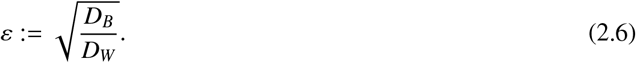

Note that decoupling the autotoxicity equation by setting ℋ = 0 is equivalent to taking the toxicity sensitivity *s* = 0.

The original KGS model can be recovered by setting ℋ = 0 in (2.4), thereby decoupling the toxicity variable *S* from the water-biomass variable pair (*U, V*). For the decoupled system, there is extensive literature on the existence and dynamics of several types of patterns, both in one and two spatial dimensions (e.g. [2, 7, 12, 13, 28, 29, 41, 48, 49, 50, 54]). In particular, the existence and dynamics of pulse solutions in the original KGS model has been studied in great detail, using a variety of techniques [12, 13, 28, 29, 41, 49, 50]. In this paper, we adopt the analytical approach as outlined in [14]. Moreover, model (2.4) has been rescaled to enable parameter identification with the original KGS model as analysed in [13]. We want to emphasise that this does not imply our model formulation is in any way preferable over alternative formulations, such as those used in [28, 29] or [49, 50]; our choice merely reflects the similarity to the method employed in [13, 48], to enable easy comparison with previous results.

## 3. Numerics

In this section we illustrate the results obtained by performing numerical simulations of equations (2.1) and (2.4) on a two-dimensional and one-dimensional spatial domain, respectively. The main aim of this section is to highlight the differences caused by the presence of toxicity when compared to ‘classical’ biomass-water models without negative plant-soil feedback. When such negative plant-soil feedback is particularly strong (i.e. for low precipitation rate, high sensitivity to toxicity, and low toxicity decay rate), two effects arise, which are not detected in biomass-water models without toxicity: the emergence of spatio-temporal (dynamic) patterns and an asymmetric biomass distribution within the patterns.

### 3.1. Simulations on a two-dimensional domain

We consider the dimensional biomass-water-toxicity model (2.1) on a two-dimensional domain Ω = {0 ≤ *x* ≤ *L*_*x*_, 0 ≤ *y* ≤ *L*_*y*_} with boundary and initial conditions

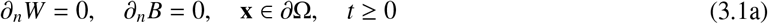

and

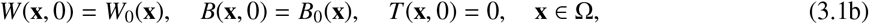

respectively. Here, ∂Ω is the boundary of Ω, ∂_*n*_ is the normal derivative on ∂Ω, *B*_0_ and *W*_0_ are initial spatial distributions of biomass and water, respectively. Our simulations are based on a numerical scheme which uses finite differences in space, and forward Euler in time. The two-dimensional domain Ω is discretised via a square lattice of 100 × 100 elements, with Δ*x* = Δ*y* = 1 and *L*_*x*_ = *L*_*y*_ = 99 meters. The initial datum for the biomass *B*_0_(**x**) satisfies *B*_0_ = 0.2 in *N*_0_ = 5000 randomly selected elements (total initial biomass 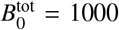) and *B*_0_ = 0 in the remaining nodes, while *W*_0_(**x**) = 40 uniformly. The simulation time *t*_*max*_ = 1000 days consists of 10^5^ time-steps Δ*t*, with Δ*t* = 0.01 days.

Since our goal is to investigate the difference between the case without toxicity (*s* = 0) and the one with strong influence of toxicity (*s* ≠ 0, small *k*), we fix all other parameter values in (2.1) as follows

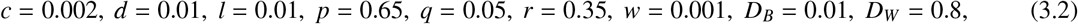

and consider two particularly representative cases for *s* and *k* (see Figure 1). In the case without toxicity (*s* = 0, Figure 1(a)), numerical simulations show the emergence of a stable stationary spatial pattern in the form of spots, consistent with existing literature results (see e.g. [26]). On the other hand, for *s* = 0.3 and *k* = 0.01, strong negative feedback due to plant toxicity induces the formation of dynamic patterns that continuously evolve in time with a propagation speed of about 3 meter/year. Moreover, the biomass distribution within the pattern is not symmetric, yielding crescent moon shapes (see Figure 1 (b)). These crescent moon shapes are observed to travel with constant velocity in the direction of highest local biomass concentration. Moreover, the velocity distribution is seen to be isotropic. Such non-symmetric patterns are usually linked to the presence of an advection term in the water equation, which models water transport due to slope (see e.g. [52, Figure 11]). Negative feedback due to autotoxicity is hence able to reproduce such non-symmetric shapes on flat terrain, in agreement with experimental observations (see [35]).

**Figure 1:**
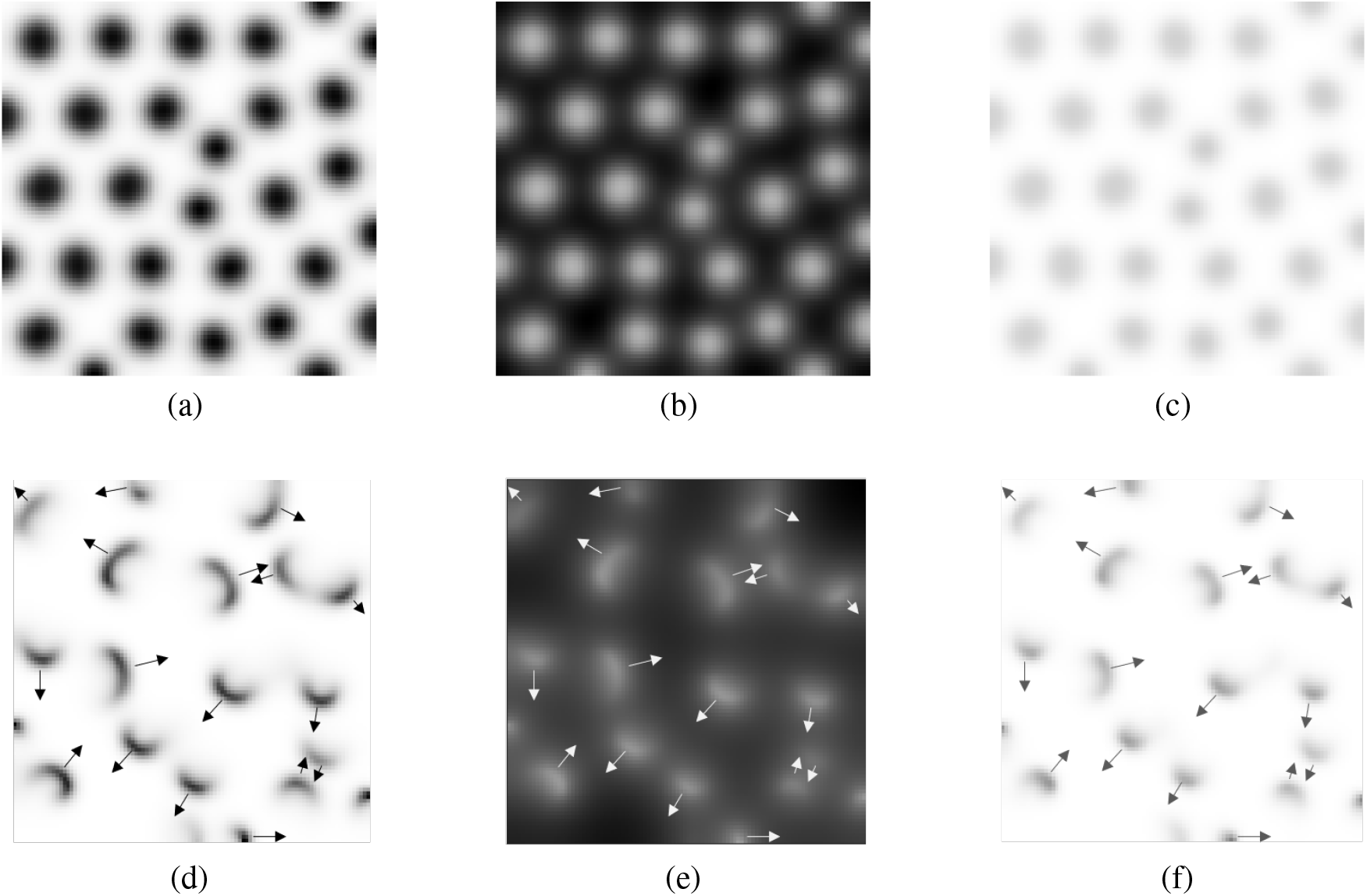
Biomass patterns obtained simulating equations (2.1) with parameter values (3.2), initial conditions and boundary conditions as in (3.1) at *t* = *t*_*max*_. Darker shades of gray correspond to higher density of *B*(**x**, *t*_*max*_), *W*(**x**, *t*_*max*_), and *T* (**x**, *t*_*max*_). The first row corresponds to simulations with *s* = 0, any *k* for (a) *B*, (b) *W*, and (c) *T*. In this case, the influence of toxicity is absent (as equations (2.1b) and (2.1c) are decoupled), and we recover (away from the boundaries) the symmetric stationary spots present in ‘classical’ biomass-water models (see [26, 27]). The second row corresponds to simulations with *s* = 0.3, *k* = 0.01 for (d) *B*, (e) *W*, and (f) *T*. Toxicity induces both a non-symmetric distribution of biomass within the spots, and movement of the spots (indicated by the arrows) with constant velocity – that is, the formation of a dynamic spatio-temporal pattern. Movies illustrating the above dynamics are available in the supplementary material of [35].

### 3.2. Simulations on a one-dimensional domain

We consider equations the nondimensionalised, rescaled model (2.4) on a one-dimensional domain 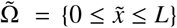 with boundary and initial conditions

**Table 2:** An overview of numerical simulation results of system (2.4) for several choices of parameters 𝒜, ℬ and 𝒟. Throughout, ℋ= 0.5 and *ε* = 0.1. () In the case 𝒜 = ℬ = 0.2, 𝒟 = 0.04, the pulses seem to be slowing down over the simulated time interval and its travelling speed tends to zero when extending the simulation to *t* = 100000, leading to a stationary pulse.

| | $\mathcal{D} = 0.04$ | $\mathcal{D} = 0.4$ | $\mathcal{D} = 4.5$ |
| --- | --- | --- | --- |
| $\mathcal{A}, \mathcal{B} = 0.02$ | Bare soil; see Figure 6 | Bare soil | Bare soil |
| $\mathcal{A}, \mathcal{B} = 0.2$ | (*) Travelling pulse(s), approximate speed 0.0042 | Stationary pulse(s) | Travelling pulse(s), approximate speed 0.0384; see Figure 3 |
| $\mathcal{A}, \mathcal{B} = 2$ | Bare soil | Bare soil | Bare soil |
| $\mathcal{A} \neq \mathcal{B}$ | Bare soil | Bare soil | Bare soil |

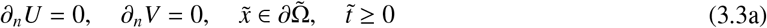

and

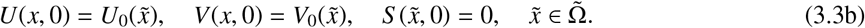

Our numerical simulations are again based on a numerical scheme which uses finite differences in space, and forward Euler in time. We consider 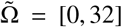 and discretize the domain by fixing *m* = 801 nodes at equal·distance 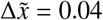. The simulation time 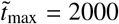 consists of 10^7^ time steps 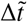, with 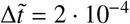.

For ℋ = 0, system (2.4) reduces to the classical KGS model. In one spatial dimension, the KGS model has been extensively studied, revealing the formation of self-replicating patterns (see e.g. [13, 28, 43]). In particular, in [13], the non-existence of travelling solitary pulses and the occurrence of pulse splitting leading to a stationary periodic pulse pattern have been shown, both analytically and numerically. In this section, our aim is to compare the numerical results presented in [13] for the toxicity-free scenario, with the scenario where (2.4b) and (2.4c) are coupled (i.e., ℋ = 0 versus ℋ ≠ 0). To this end, we fix all parameters except for ℋ as follows:

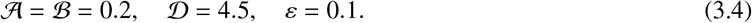

These values are commensurate with the parameter choices in the two-dimensional simulations (3.2), with in addition *k* = 0.01, and *s* = 0 (no toxicity sensitivity) or *s* = 0.3 (high toxicity sensitivity). As initial data, we take 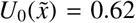 uniformly, which is commensurate with the choice *W*_0_(**x**) = 40 made in subsection 3.1. For *V*, we consider the following two initial profiles:

- a Gaussian peak in the centre of the domain

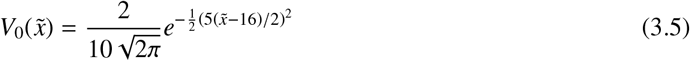
- a “half-bare–half-vegetated” domain

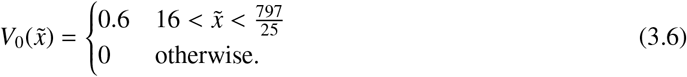

(We note that the condition *V*_0_ = 0 at 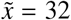 is motivated by our intent to analyse the effect of different boundary conditions while keeping the same initial configuration for *V*.) When ℋ = 0 (see Figure 2), we observe the formation of a stationary, periodic, spotted pattern as a consequence of pulse splitting, independent of the choice of the initial *V*-profile. This observation is in agreement with the results presented in [13, 26] and with the two-dimensional numerical simulations presented in subsection 3.1. When we now ‘turn on’ the influence of toxicity by choosing ℋ = 0.5, the numerical simulations show the emergence of travelling pulses. In particular, when the initial profile 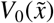 is chosen as in (3.5), the initial pulse symmetrically splits into two pulses which periodically move away from each other and toward each other within the domain (see Figure 3(a)). On the other hand, if we take the “half-vegetated–half-bare” initial profile (3.6), we observe that a single pulse forms, which periodically travels back and forth within the domain (see Figure 3(b)). In both cases, numerical results reveal that the travelling pulse is asymmetric (see Figure 4) and that the speed of such pulses is approximately 0.0384, which could be interpreted 𝒪(*ε*^2^) for this choice of *ε*. This value is of the same order of magnitude with respect to the numerical propagation speed computed for the 2D simulations.

**Figure 2:**
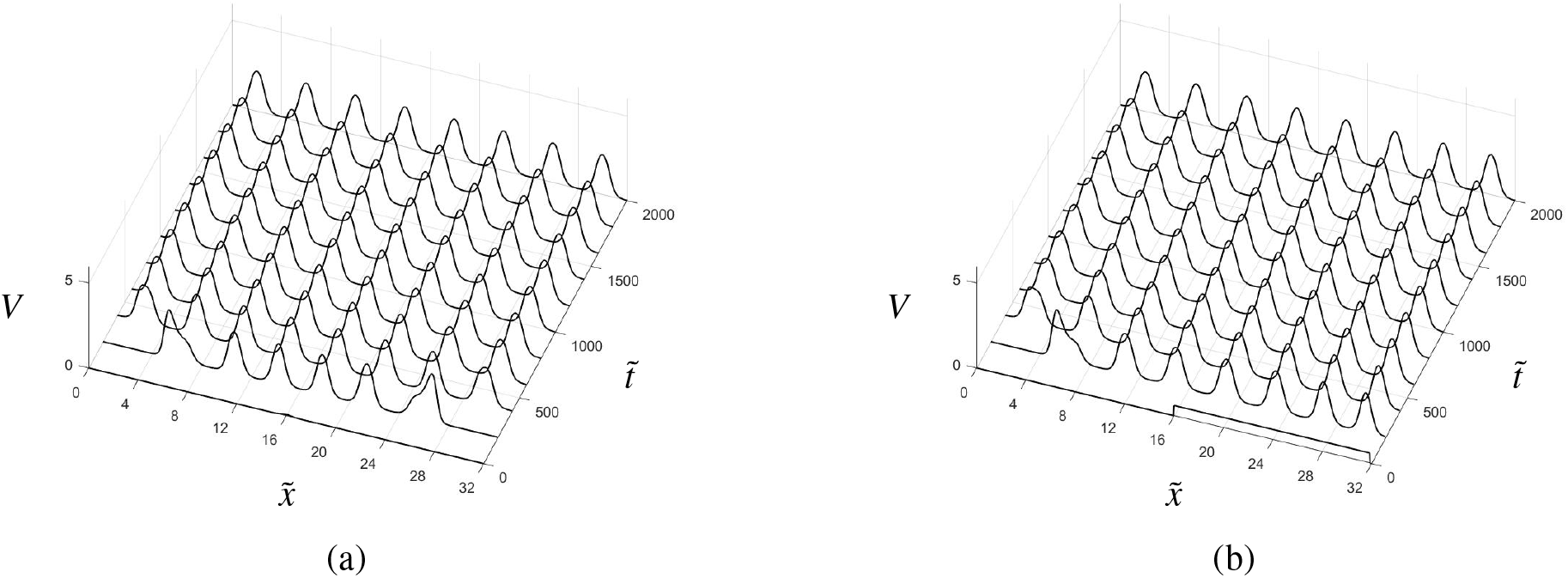
Evolution of *V* in space 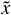 and time 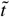 obtained by numerically simulating equations (2.4) with parameters as in (3.4) and ℋ = 0. (a) Simulations obtained for an initial datum 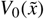 as in (3.5). (b) Here, 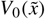 is fixed as in (3.6). In both cases, decoupling the dynamics of *V* from those of *S* leads to a symmetric, stable, regular pulse pattern through a process of pulse splitting.

**Figure 3:**
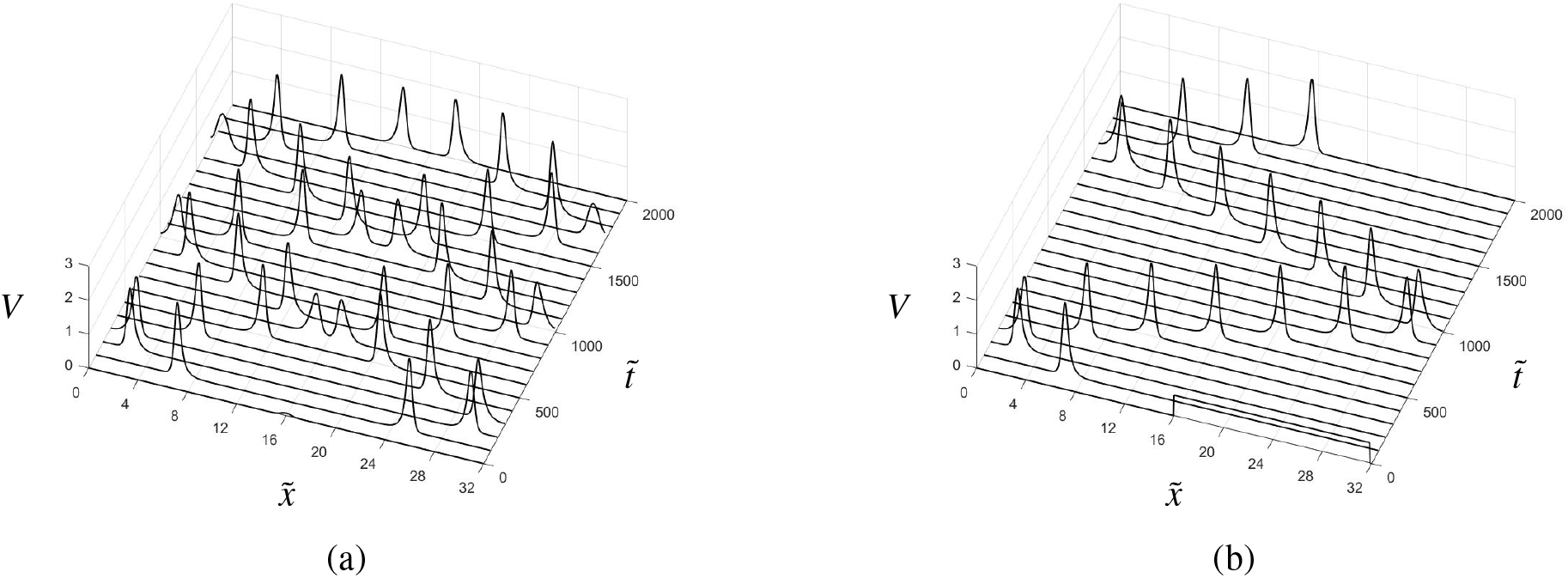
Evolution of *V* in space 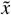 and time 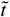 obtained by numerically simulating equations (2.4) with parameters as in (3.4) and ℋ = 0.5. (a) Simulations obtained for an initial datum 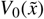 as in (3.5). The strong influence of *S* on the dynamics of *V* induces the formation of a symmetric pair of pulses, which travel within the domain, moving away from and towards each other periodically. (b) Here, 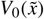 is chosen as in (3.6). In this case, a single pulse forms, which travels back and forth along the spatial domain. Note that, in both cases, *V* is not symmetrically distributed within the pulse.

**Figure 4:**
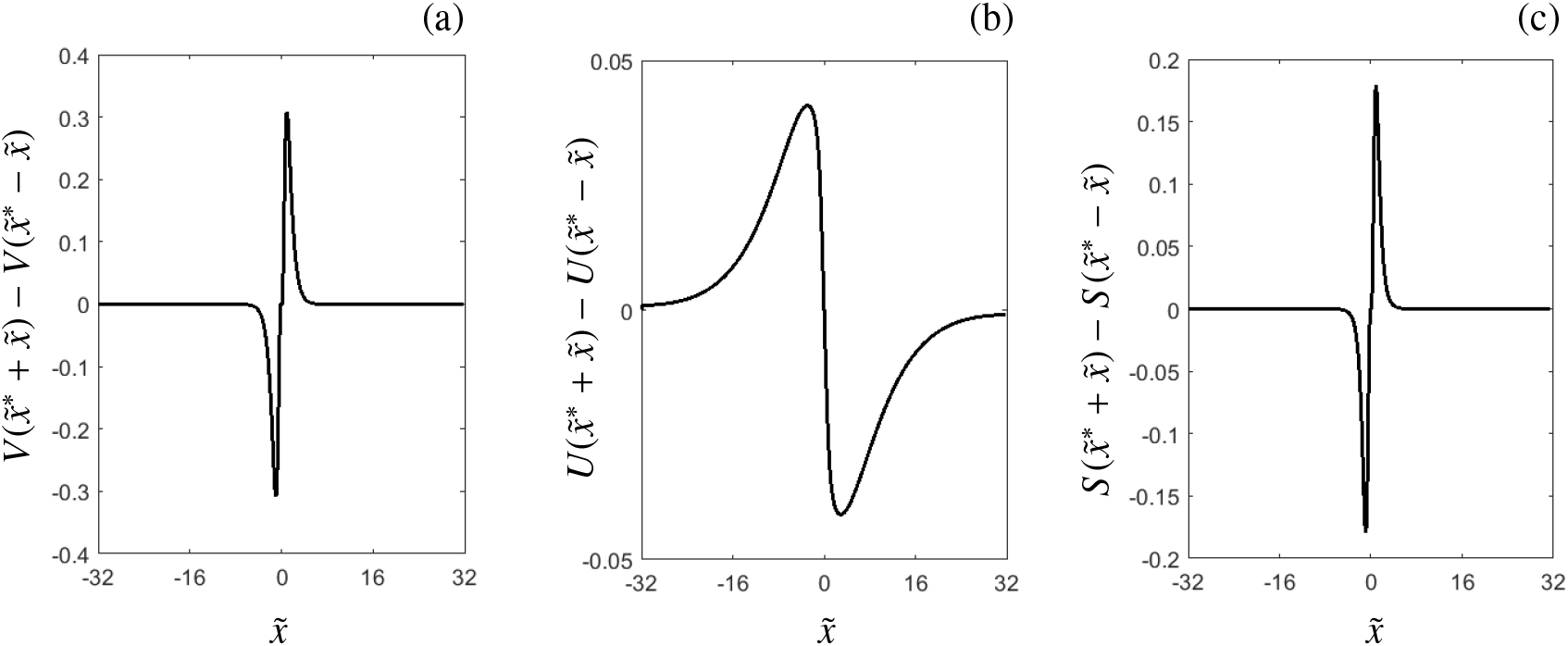
Measurement of asymmetry of the travelling pulse obtained by simulating system (2.4) until time 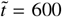, starting from 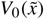 as in (3.6). The symbol 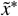 indicates the *x*-value where *V* reaches its maximum value (peak of the pulse). The panels show the asymmetry function 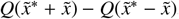, where (a) *Q* = *V*, (b) *Q* = *U*, and (c) *Q* = *S*.

A snapshot of the simulated travelling pulse is shown in Figure 5.

**Figure 5:**
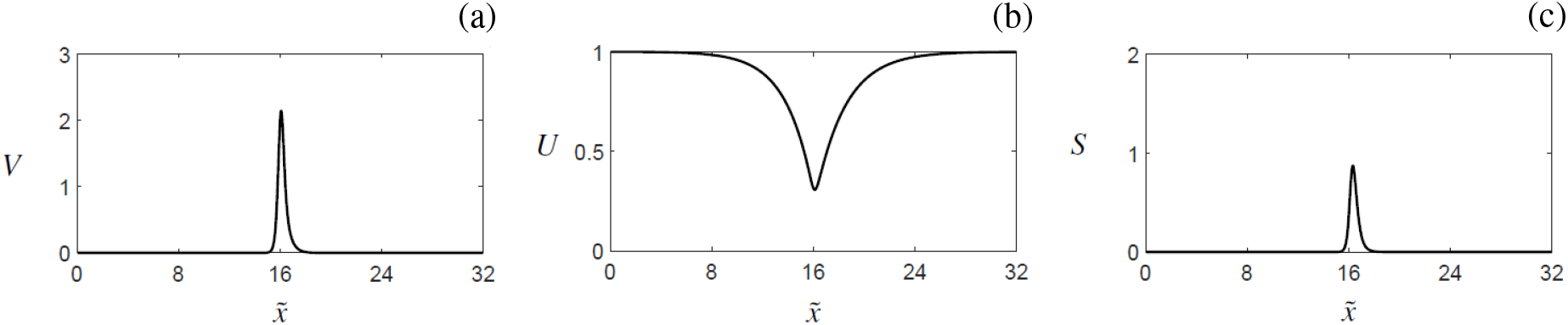
Snapshot of the a travelling pulse at time 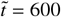 obtained by simulating system (2.4) starting from 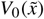 as in (3.6) with parameters as in (3.4). (a) *V*-component, (b) *U*-component, (c) *S* -component.

#### *Remark* 3.1

(*Boundary conditions*). The results presented in this section appear to be independent of boundary conditions, as the travelling pulse structure persists when considering different (larger) domains which mimick an infinite spatial interval. We are mainly interested in the dynamics occurring away from the boundaries: therefore, even though the interaction with the boundary might lead to some transient effects, (see e.g. Fig. 3), their relevance in our analysis can be neglected.

#### *Remark* 3.2

(*Simulations time*). Numerical evidence strongly suggests that the findings presented in Section 3.1 and 3.2 are not affected by the length of the simulations times, i.e. they persist when considering longer intervals mimicking long-time behaviour. Therefore, we focus our attention on finite times, with *t*_*max*_ = 1000 days for 2D simulations and 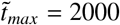 in the case of 1D simulations.

Remarks 3.1 and 3.2 suggest that the travelling pulse observed in the simulations is stable in the PDE sense.

A preliminary, order-of-magnitude investigation into the role of the parameters 𝒜, ℬ and 𝒟 is presented in Table 2. An important observation is that the existence of pulse solutions not only clearly depends on the magnitude of 𝒜 and ℬ, but also on the fact that 𝒜 and ℬ have to be chosen (approximately) equal. Secondly, by varying 𝒟, we are able to find stationary pulse solutions. In addition, for relatively low values of𝒜, ℬ and 𝒟, simulations show strongly asymmetric, travelling pulse solutions –or travelling fronts with a decaying back– as a transient phase towards bare soil, see Figure 6. To our knowledge, transients of this shape have not been observed in the context of the classical KGS model.

**Figure 6:**
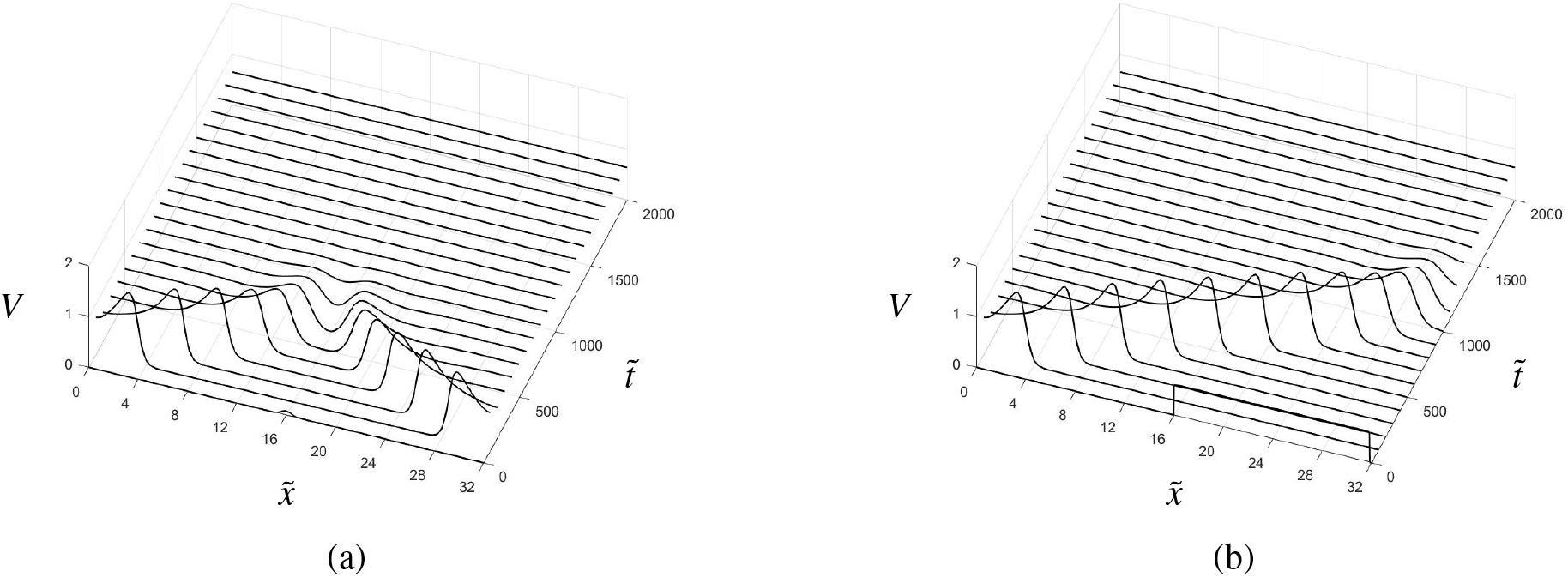
Evolution of *V* in space 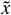 and time 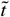 obtained by numerically simulating equations (2.4) with 𝒜= ℬ= 0.02, 𝒟= 0.4, ℋ= 0.5, and *ε* = 0.1. (a) Initial datum 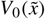 as in (3.5). Here, two travelling fronts form, meet at the centre of the domain, and then merge and decay to a bare soil state. (b) 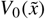 as in (3.6). In this case, a single travelling front emerges, and eventually decays to the bare soil state as well.

## 4. Existence of pulse solutions

In this section, we investigate the existence of stationary and travelling pulse solutions to system (2.4) on an unbounded, one-dimensional spatial domain 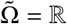. We confine our study to pulse solutions that are biasymptotic to the trivial homogeneous stable steady (bare soil) state (*U, V, W*) = (1, 0, 0). System (2.4) in one spatial dimension is restated here for completeness:

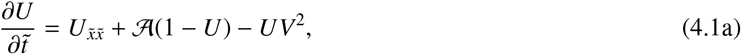

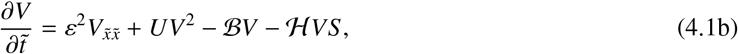

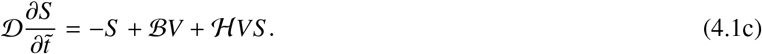

### *Remark* 4.1

(*Homogeneous steady states*). Apart from the trivial homogeneous steady state (*U, V, S*) = (1, 0, 0), system (4.1) admits two nontrivial homogeneous steady states, given by

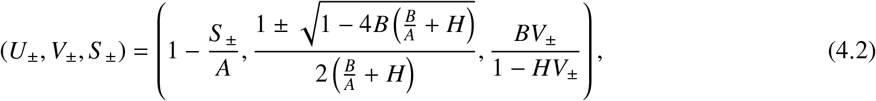

which exist for 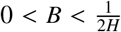 and 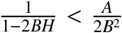. The limit *H* → 0 yields the pair of nontrivial homogeneous steady states in the original Klausmeier model, with existence condition 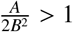. However, one can choose *A* and *B* such that the nontrivial solution pair (4.2) does not exist for *H* below (alternatively, above) a certain threshold value. Hence, the family (4.2) is not for every *A, B* and *H* a continuous extension of the pair of nontrivial steady states in the original Klausmeier model.

We introduce a co-moving frame coordinate

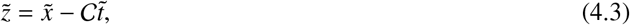

in which system (4.1) takes the form

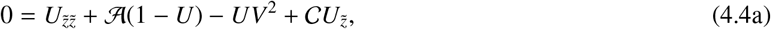

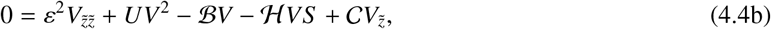

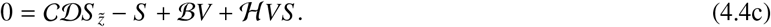

We look for pulse solutions, i.e. solutions to (4.4) that are bi-asymptotic to the trivial background state (*U, V, S*) = (1, 0, 0), that is, solutions (*U*(*z*), *V*(*z*), *S* (*z*)) for which lim_*z*→±∞_(*U*(*z*), *V*(*z*), *S* (*z*)) = (1, 0, 0). To that end, we make the crucial assumption that *ε*, which measures the biomass diffusion rate with respect to the water diffusion rate, is an asymptotically small parameter, i.e. 0 < *ε* ≪ 1. This is consistent with the experimental data collected in arid environments (see, e.g., [45] and references therein). Next, in accordance with previous work [12, 13, 48, 54], we allow all model variables and parameters to scale with (a power of) this small parameter *ε*. While the introduction of these scalings can seem unnecessary complicated and confusion due to the large number of new parameters that are introduced as scaling exponents, previous analysis [12, 13, 48, 54] has shown that several solution types –and pulse patterns in particular– exist only in regions of parameter space that scale with *ε* in a particular way; moreover, the resulting pattern amplitude may also scale with *ε*. A preparatory asymptotic scaling analysis of system (4.4), along the lines of [13, 48], can be found in AppendixA. The resulting rescaling can be summarised as follows:

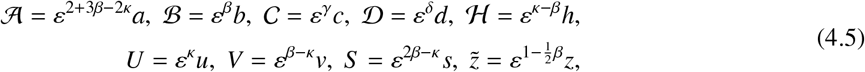

with the additional assumptions that

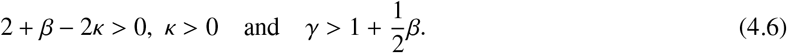

(4.6)

Note that the positivity of the original model parameters 𝒜, ℬ, 𝒟, ℋ implies that their rescaled counterparts *a, b, d, h* are positive and 𝒪 (1) in *ε*; however, the associated scaling exponents may be negative. This also applies to the model variables *U, V, S* and their rescaled counterparts *u, v, s*. The speed of the moving frame,, can take any sign.

Application of rescaling (4.5) to (4.4) yields the following 5-dimensional dynamical system:

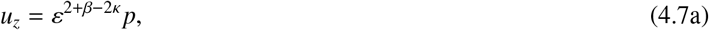

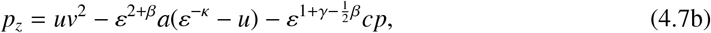

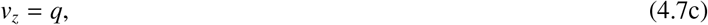

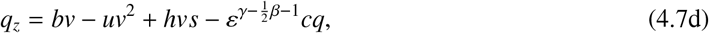

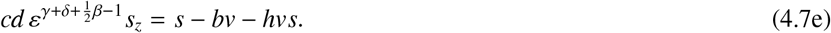

Pulse solutions to (4.4) can now be identified with orbits in system (4.7) that are homoclinic to the (rescaled) trivial background state (*ε*^−κ^, 0, 0, 0, 0). In the following, we will use techniques from geometric singular perturbation theory [12, 13, 14, 48] to constructively establish the existence of such homoclinic orbits in system (4.7).

### 4.1. Stationary pulses

For stationary pulses, we have 𝒞 = 0 in (4.3), hence *c* = 0 in (4.7). We obtain the differential-algebraic system

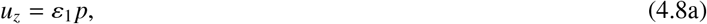

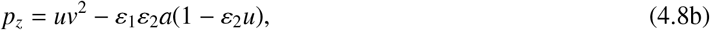

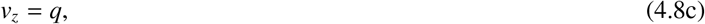

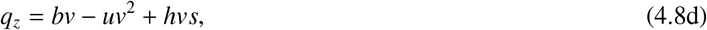

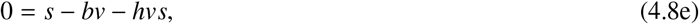

where we have defined the two small parameters

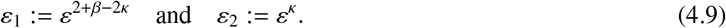

Note that *ε*_1,2_ are indeed asymptotically small by the scaling conditions (4.6). However, the case *ε*_2_ = 1 is also covered by the analysis presented in this paper, but has been omitted for presentation purposes. For more information, we refer to AppendixA, in paricular the remark below (A.10).

First, we observe that the algebraic equation (4.8e) can be solved for *s*, yielding

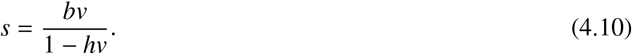

From the biological interpretation of the model, it follows that all model components are assumed to be nonnegative, which implies that

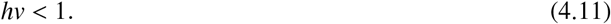

In other words, the dynamics of (4.8) take place on the 4-dimensional invariant manifold ℳ_0_ embedded in 5-dimensional (*u, p, v, q, s*)-phase space, that is given by

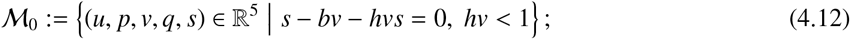

see also Figure 9 (a). At this point, the introduction of ℳ_0_ can seem somewhat superfluous; however, this manifold will also play a role in the upcoming analysis of travelling pulses (see Section 4.2), and its introduction can illuminate similarities between the analysis of the current section and that of section 4.2. On ℳ_0_, the dynamics are given by the 4-dimensional dynamical system

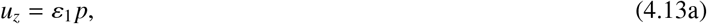

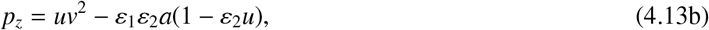

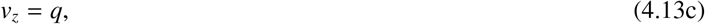

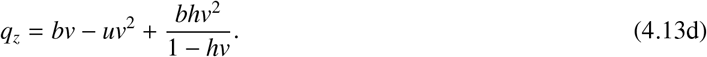

Note that, for *h* = 0, this is exactly the system studied in [12, 13]. For future reference, we introduce the 2-dimensional hyperplane

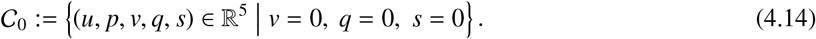

We note that 𝒞_0_ is invariant under the flow of (4.8); moreover, 𝒞_0_ ⊂ ℳ_0_ (4.12).

#### *Remark* 4.2

(*Fast and slow variables*). In system (4.13), the *p*-variable cannot a priori be identified as ‘slow’ due to the presence of the *uv*^2^-term in (4.13b). However, this does not preclude the application of geometric singular perturbation theory; specifically, from the analysis presented below and in previous work [12, 13, 14, 47, 48], it follows that the *p*-component expresses both slow and fast behaviour.

#### 4.1.1. Fast dynamics

Following the approach of geometric singular perturbation theory, we study the fast reduced limit of (4.13) by letting *ε*_1_ → 0, and obtain the planar, Hamiltonian, fast reduced system

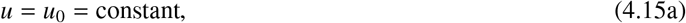

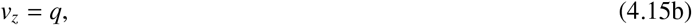

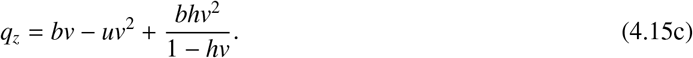

The dynamics of *p* can be obtained by direct integration of *p*_*z*_ = *u*_0_*v*^2^. Note that 𝒞_0_ (4.14) consists of hyperbolic (trivial) equilibria of the fast reduced system (4.15); hence, 𝒞_0_ is a normally hyperbolic invariant manifold [30]. We introduce

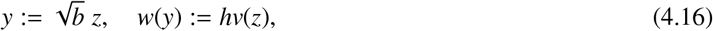

to obtain

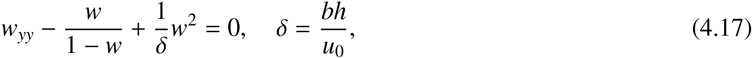

with Hamiltonian

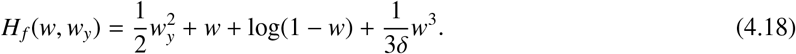

Analogously to [12, 13, 14, 48], we look for a homoclinic orbit to the origin in system (4.17); note that this orbit lies on the level set *H*_*f*_ = 0. Straightforward phase plane analysis reveals that such a homoclinic orbit exists as long as 0 < *δ* < *δ*_max_, and the maximally attained *w*-value of the associated ‘spike’ solution is given by the unique positive solution to

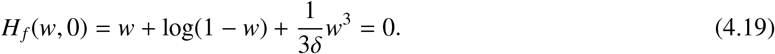

The value for *δ*_max_ can now be determined by considering the situation when (4.19) is degenerate, which is when *δ* = *w*(1 − *w*). This degeneracy occurs when the graph of *H*_*f*_ (*w*, 0) is tangent to the *w*-axis for the unique positive solution to (4.19), causing the homoclinic orbit to deteriorate in a pair of heteroclinic orbits, as shown in Figure 7 (b). Hence, we find that

**Figure 7:**
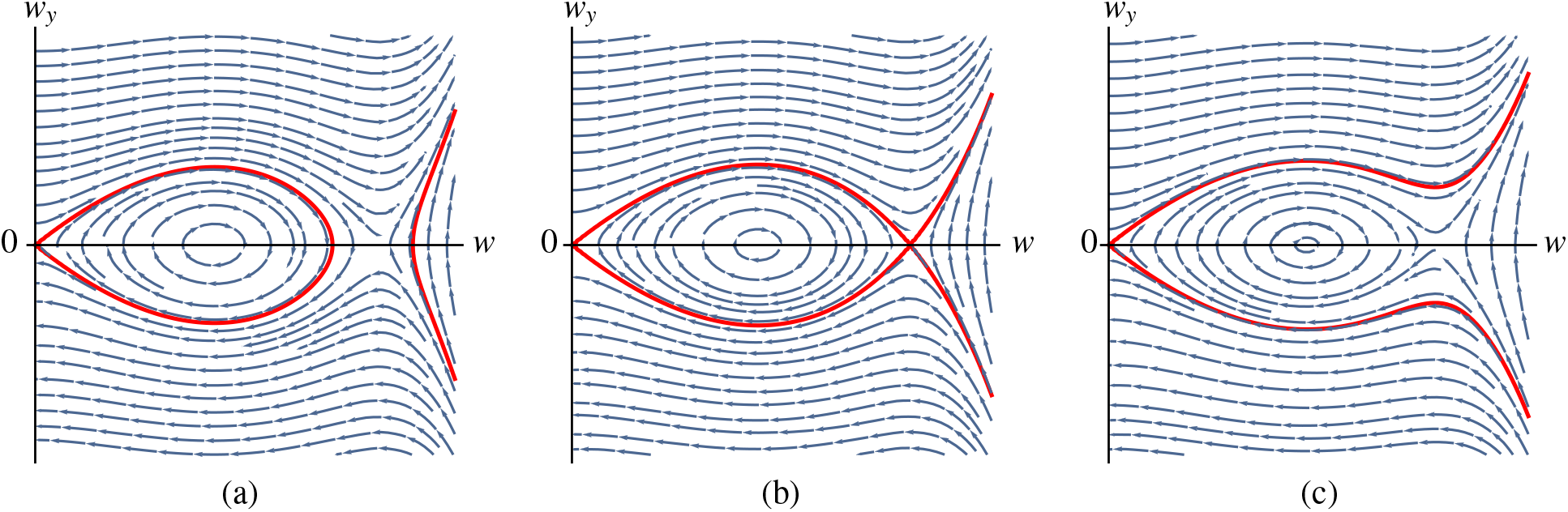
The phase plane of (4.17) for varying values of *δ*. The level set *H*_*f*_ = 0 is indicated in red. (a) *δ* = 0.225 < *δ*_max_, a planar homoclinic orbit exists. (b) *δ* = *δ*_max_ (4.22), the planar homoclinic orbit deteriorates in a pair of heteroclinic orbits. (c) *δ* = 0.235 > *δ*_max_, no planar homoclinic orbit to the origin exists.

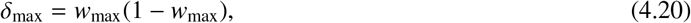

where *w*_max_ is the unique positive solution to

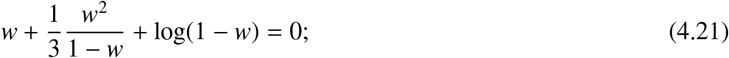

the approximate numerical values are

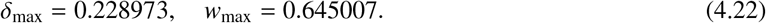

See Figure 7 for an illustration of the dynamics of (4.17).

Note that, although the unique homoclinic solution *w*_*h*_(*y*; *δ*) to (4.17) does not have a closed-form expression, it can be approximated by substitution of an asymptotic expansion in *δ*, as 0 < *δ* < *δ*_max_ ≈ 0.23 (4.22). Writing

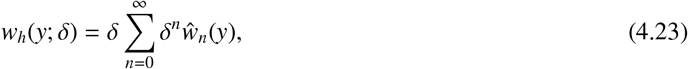

we find for the first terms

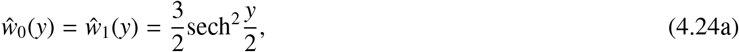

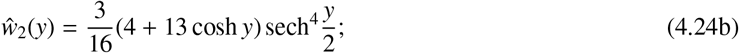

see Figure 8 for a visualisation of the approximation accuracy of series (4.23).

**Figure 8:**
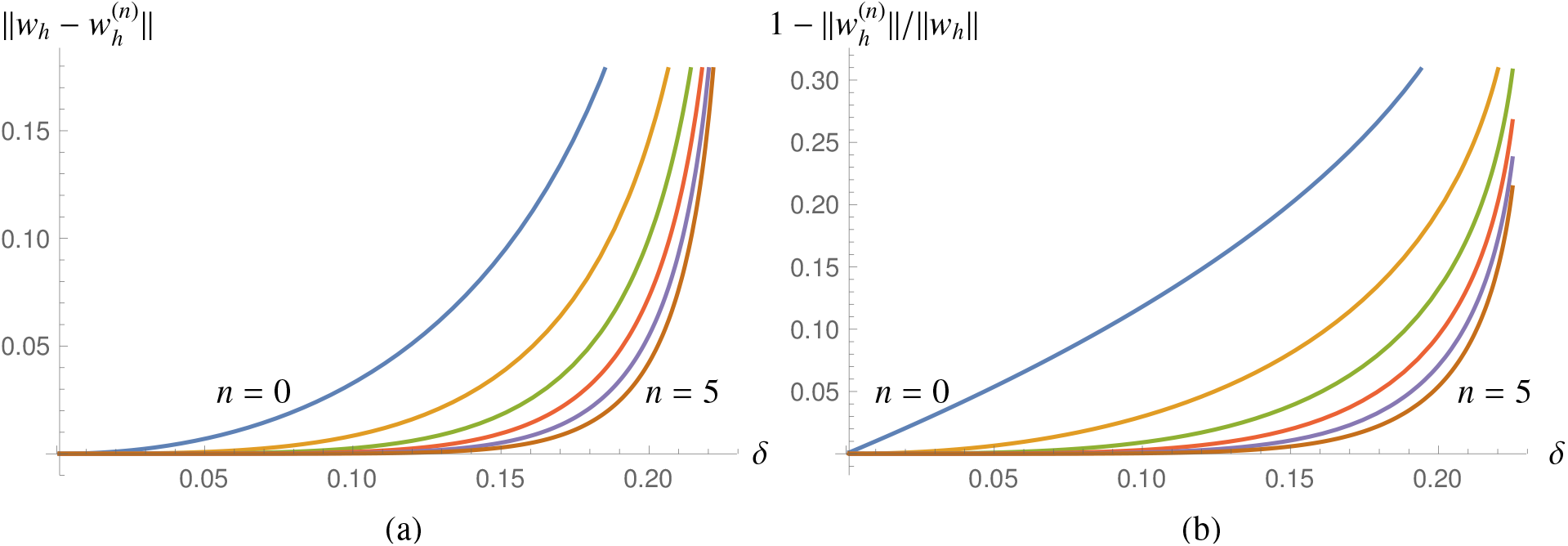
The approximation accuracy of the asymptotic series (4.23). (a) The *L*^2^-norm of the difference of *w*_*h*_(*y*; *δ*), the unique homoclinic solution to (4.17) (computed numerically), and its *n*-th order asymptotic approximation 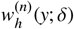 according to (4.23). (b) The relative error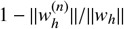.

**Figure 9:**
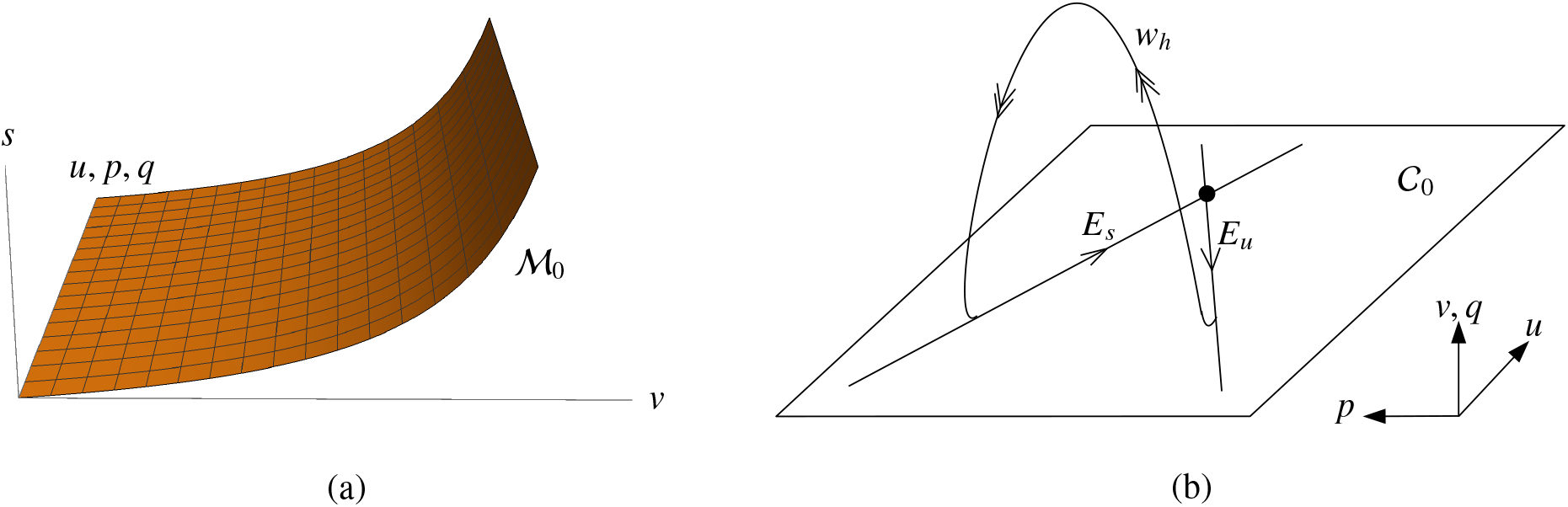
The geometric construction of the stationary pulse solution underlying Theorem 4.3. (a) The 4-dimensional invariant manifold ℳ_0_ (4.12) embedded in 5-dimensional (*u, p, v, q, s*)-phase space. (b) On ℳ_0_, the slow (*u, p*)-dynamics take place on the invariant manifold _0_ (4.14). The fast dynamics normal to ℳ_0_, can be used to connect the unstable and stable subspaces *E*_*u*_ and *E*_*s*_ (4.26) through the fast homoclinic orbit *w*_*h*_ (see subsection 4.1.1); see also [13, Figures 2 and 3].

#### 4.1.2. Slow dynamics

As noted at its definition, 𝒞_0_ (4.14) is a normally hyperbolic manifold that is invariant under the flow of (4.8). The flow on 𝒞_0_ is given by

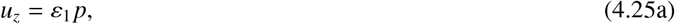

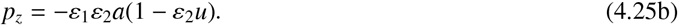

This flow is linear; it has a unique saddle equilibrium at 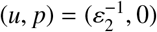, with stable and unstable manifolds given by the stable and unstable linear subspaces

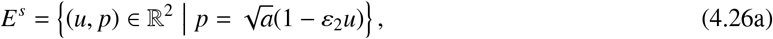

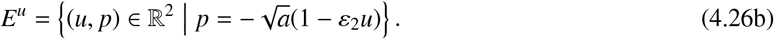

Note that the dynamics of (4.25) are slow in *z*; the eigenvalues of the saddle equilibrium are of order O(*ε*_1_*ε*_2_).

#### 4.1.3. Constructing a stationary pulse solution

We establish the existence of a stationary pulse solution to (4.1) by constructing a homoclinic orbit in system (4.8). This construction, which uses techniques and concepts from geometric singular perturbation theory, is based directly on the equivalent construction of pulses in the classical KGS model, as carried out in e.g. [13]. Due to the abundance of high quality sources, we choose to highlight central concepts in the pulse construction below. For detailed arguments and proofs, we refer to [12, 13, 14, 48].

The pulse solution that we want to construct, is an orbit that is homoclinic to the equilibrium 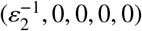 in (4.8). This equilibrium lies in particular on the invariant manifold 𝒞_0_ (4.14). In order to construct a homoclinic orbit, we need to consider the flow both on 𝒞_0_ and normal to 𝒞_0_. The latter can be studied from a geometric viewpoint by considering the unstable and stable submanifolds of 𝒞_0_ under the flow (4.8), denoted by *W*^*u,s*^(𝒞_0_), respectively. To leading order in *ε* (and hence to leading order in *ε*_1,2_), the flow normal to 𝒞_0_ is given by the fast reduced system (4.15), as studied in subsection 4.1.1. The existence of a homoclinic orbit in this planar system implies that *W*^*u*^(𝒞_0_) and *W*^*s*^(𝒞_0_) intersect. Moreover, using the reversibility symmetry of (4.8) (i.e. the invariance of the flow (4.8) of under the reflection (*z*; *p, q*) → (−*z*, −*p*, −*q*)), one can show that this intersection is transversal; for details, see [13, 48]. Using the homoclinic solution *w*_*h*_(*y*; *δ*) to (4.17), we can track the fast flow normal to 𝒞_0_ through this intersection. We define the interval

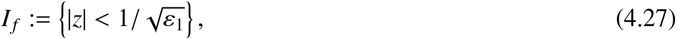

chosen such that *I* _*f*_ is asymptotically large in *z* but asymptotically small in *ε*_1_*z*. We calculate the change of the slow variables *u* and *p* through the fast flow, over this interval, as follows. From (4.8), we see that *u*_*z*_ is 𝒪 (*ε*_1_), hence *u* is constant to leading order in *z*; we write

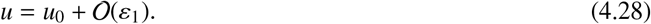

Moreover, *p*_*z*_ is to leading order slaved to *u* and *v*; we write

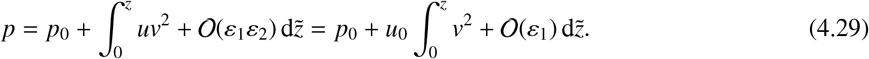

We observe that, to leading order, *p*− *p*_0_ is odd in *z*. The change of *u* and *p* over *I* _*f*_ can now be calculated to leading order as

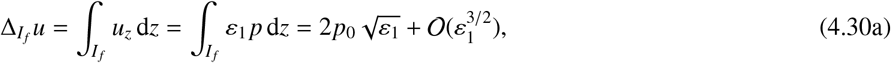

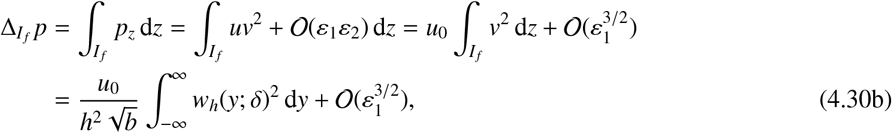

cf. (4.16). Geometric singular perturbation theory [13, 14, 22, 30] enables us to construct an orbit by concatenating orbits on 𝒞_0_ and normal to 𝒞_0_; in particular, any orbit homoclinic to 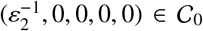 first flows away from this equilibrium along (and exponentially close to) *E*^*u*^ (4.26). Then, it takes an excursion away from 𝒞_0_, during which its evolution is to leading order determined by the fast reduced flow (4.15). After touching down again exponentially close to C_0_, the *u*-component has not changed its value to leading order, but the *p*-component has, cf. (4.30). For the orbit to be biasymptotic to the equilibrium 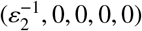, this change needs to be such that the touchdown point lies exponentially close to *E*^*s*^ (4.26), such that the last, slow, orbit component takes us back to the 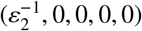 along *E*^*s*^; see Figure 9 for a sketch of the geometric situation and construction.

For this concatenation construction to work, we find the condition that the ‘vertical’ *p*-distance between *E*^*u*^ and *E*^*s*^ must be equal to the leading order change in *p* during the fast excursion over the interval *I* _*f*_, as calculated in (4.30), that is,

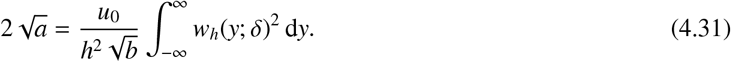

Using techniques from geometric singular perturbation theory, one can show that, for sufficiently small *ε*_1_, condition (4.31) is not only *necessary*, but also *sufficient* to prove the existence of a homoclinic orbit in (4.8) that is asymptotically close to its singular concatenation. This existence follows from the persistence of 𝒞_0_ and its stable and unstable manifolds *W*^*u,s*^(𝒞_0_), together with the observation that *W*^*u*^(𝒞_0_) and *W*^*s*^(𝒞_0_) intersect transversally; for more details and proofs, see [13, 14, 48]. For the purposes of this paper, it is sufficient to consider the outcome of previous equivalent analyses, namely that stationary pulse solutions are completely characterised by the existence condition (4.31), in the following way.

##### Theorem 4.3.

*Let ε*_1_ *be sufficiently small, and let u*_*_ *be a nondegenerate solution to*

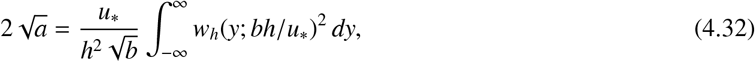

*where w*_*h*_(*y*; *bh*/*u*_*_) *is the unique, positive, nontrivial solution to* (4.17) *for which* lim_*y*±∞_ *w*_*h*_(*y*; *bh*/*u*_*_) = 0. *Then, system* (4.8) *admits an orbit that is homoclinic to the equilibrium* 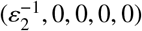. *To leading order in ε*_1_, *this orbit is given by*

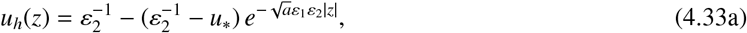

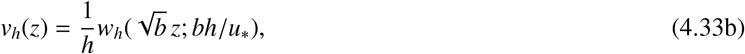

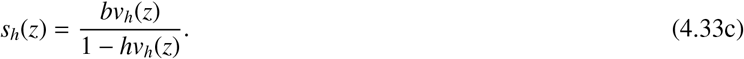

*Conversely, if* (4.32) *has no solution, then no orbit homoclinic to* 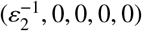 *exists*.

*Proof*. The proof is completely analogous to that of [13, Theorem 4.1], with the fast planar homoclinic *w*_*h*_(*y*; *δ*) taking the role of *v*_0_ [13, equation (3.3)] therein. Since the arguments of the proof of [13, Theorem 4.1] depend only on the *existence* of such a planar homoclinic orbit, and not on its specific functional form, the reasoning in [13] holds ad verbatim for the construction described in section 4.1 which leads to the statement of Theorem 4.3. □

##### *Remark* 4.4

(*Role of ε*_2_). Although (4.8) contains two asymptotically small, independent parameters *ε*_1_ and *ε*_2_, only *ε*_1_ needs to be sufficiently small. The reason for this is that the time scale difference in (4.8) is only determined by *ε*_1_ (4.27), and that the correction terms for the leading order expressions used in the geometric construction (e.g. (4.30)) only depend on *ε*_1_. The small parameter *ε*_2_ solely measures the *u*-value of the homogeneous background state 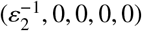. Indeed, the analysis leading to Theorem 4.3 applies ad verbatim for the case *ε*_2_ = 1; see also AppendixA.

Note that condition (4.31) can be formulated in terms of *δ* As

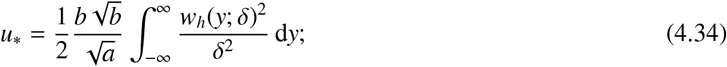

taking the limit *h* → 0, i.e. *δ* → 0 (4.17), yields

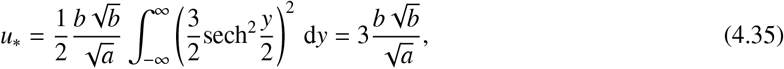

cf. (4.23) and (4.24). This is in accordance with the observation that the limit *h* → 0 of system (4.8) yields the classical KGS system, for which (4.35) has been derived as the existence condition of stationary pulse solutions [13, equation (4.2)]. See Figure 10 for the behaviour of *u*_*_ for increasing values of *h*, and for a plot of the pulse solution given in Theorem 4.3.

**Figure 10:**
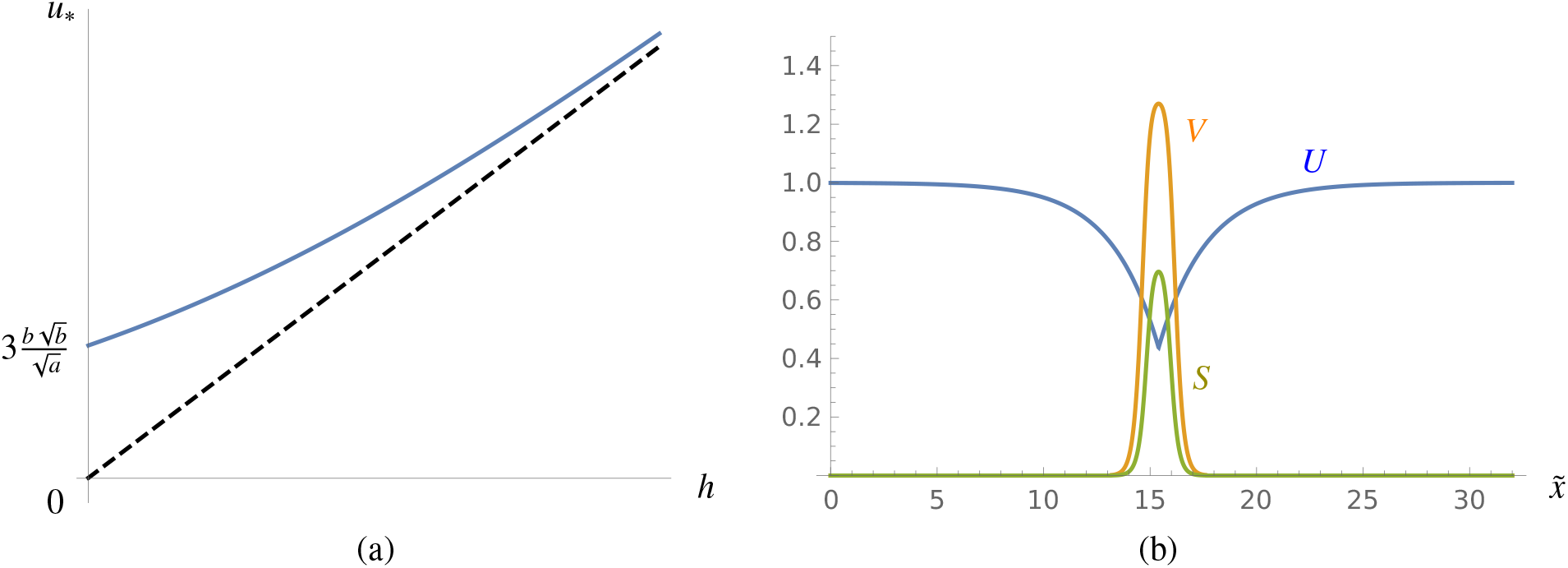
(a) The value of *u*_*_, solving the existence condition (4.32) for increasing values of *h* (with *a* and *b* fixed), in blue. The set of admissible values, for which 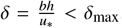, is bounded below by the dashed line 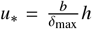. (b) A plot of the pulse solution established in Theorem 4.3, for the parameter values (3.4), with *U* in blue, *V* in orange and *S* in green.

### 4.2. Travelling pulses

Pulses travelling with nonzero speed 𝒞≠ 0, i.e *c* ≠ 0, are homoclinic orbits to system (4.7). In contrast to the stationary case studied in section 4.1, the dynamics of this system are fully 5-dimensional. For clarity of presentation, we introduce the small parameter

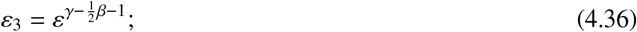

note that *ε*_3_ is indeed asymptotically small by conditions (4.6). Using the previously defined small parameters *ε*_1,2_ (4.9), we can rewrite system (4.7) as

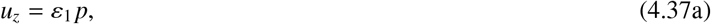

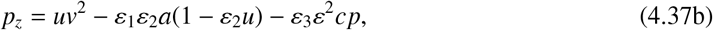

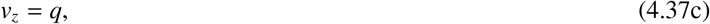

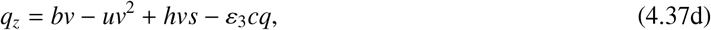

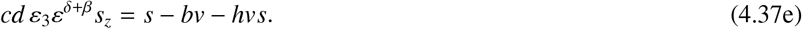

This system can be treated in a similar fashion as in section 4.1. However, there are important differences. First, we observe that, unlike in the stationary situation (see section 4.1.1), the time scale of the dynamics of (*v, q, s*) in system (4.37) is not immediately clear, because the asymptotic magnitude of *ε*^*δ*+*β*^ is not yet determined. In particular, depending on the sign and magnitude of *δ* + *β*, the dynamics of *s* may be faster, equivalent, or slower than those of (*v, q*). We examine all three cases, and the subsequent construction of a travelling pulse, in subsections 4.2.2, 4.2.3 and 4.2.4. Secondly, it is important to note that that the invariant 2-dimensional hyperplane 𝒞 _0_ (4.14) that was defined in the context of stationary pulses, is also invariant under the flow of the ‘nonzero speed system’ (4.37). The dynamics of (4.37) on 𝒞 _0_ will be analysed in subsection 4.2.1.

#### 4.2.1. Slow dynamics

On the invariant hyperplane 𝒞 _0_ (4.14), the dynamics of (4.37) are determined by the planar system

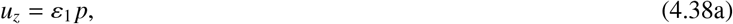

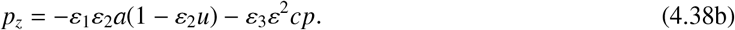

As in the stationary case (see subsection 4.1.2), the dynamics on 𝒞_0_ are linear. Again, the only equilibrium is the saddle at 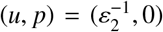. However, due to the advection term, the flow is not symmetric anymore with respect to reflection in the *u*-axis (compare (4.25) and (4.26)); instead, the stable resp. unstable manifolds of the saddle equilibrium are given by the stable resp. unstable linear subspaces

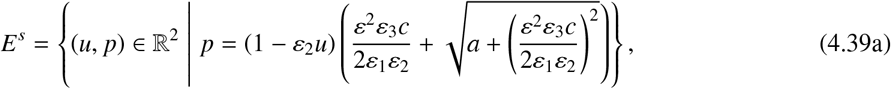

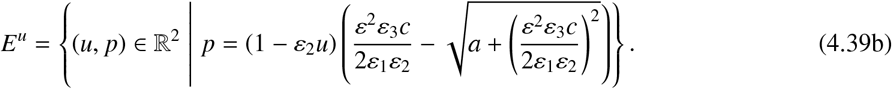

The dynamics of (4.38) are slow in *z*, as the eigenvalues of the saddle equilibrium are of order 𝒪 (max(*ε*_1_*ε*_2_, *ε*_3_*ε*^2^)) – that is, asymptotically small.

#### 4.2.2. Constructing a travelling pulse solution, case I: s faster than (v, q)

We assume that the *s*-dynamics are faster than the (*v, q*)-dynamics, that is, *ε*_3_*ε*^*δ*+*β*^ → 0 as *ε* → 0. Taking the limit *ε* → 0 in (4.37) then yields

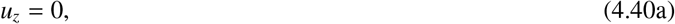

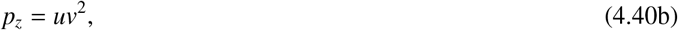

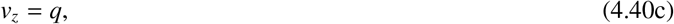

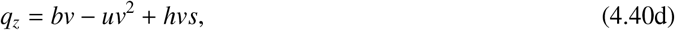

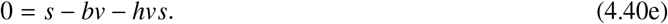

This is precisely the same system as studied in subsection 4.1.1. In particular, the same algebraic equation as (4.8e) defines the same 4-dimensional manifold ℳ _0_ (4.12). Introducing the fast coordinate

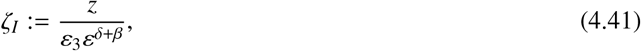

system (4.37) takes the form

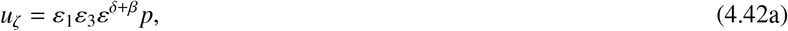

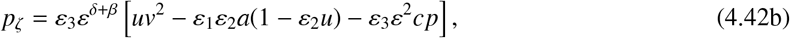

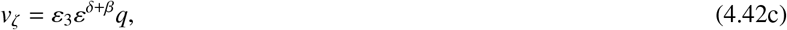

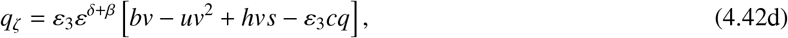

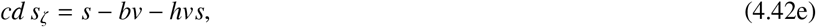

with singular limit

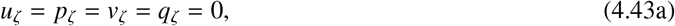

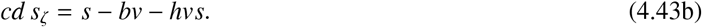

It is easy to check using (4.43) that ℳ_0_, which consists entirely of equilibria of the reduced fast system (4.43), is normally hyperbolic. Hence, by geometric singular perturbation theory (see e.g. [22, 30]), for sufficiently small *ε*, there exist a normally hyperbolic manifold ℳ _*ε*_ that is invariant under the flow of the full 5-dimensional system (4.37); moreover, ℳ_*ε*_ is O(*ε*_3_*ε*^*δ*+*β*^) close to ℳ_0_.

In contrast to the stationary case (4.8), the dynamical system we are investigating here (4.42) is fully 5-dimensional; hence, we need to look at the dynamics normal to ℳ_*ε*_. Since these normal dynamics are one-dimensional, ℳ_*ε*_ is either uniformly attracting or uniformly repelling, depending on the sign of *c*. In either case, it follows that any *bounded* (in particular, any homoclinic) orbit of (4.37) must lie entirely on ℳ_*ε*_. By the uniform asymptotic proximity of ℳ_*ε*_ to ℳ_0_, we can determine the dynamics on ℳ_*ε*_ by a regular perturbation expansion of system (4.37) in powers of *ε*_3_*ε*^*δ*+*β*^.

We also observe that, as _0_ is invariant under the full flow of (4.37), it necessarily holds that 𝒞_0_ ⊂ ℳ_*ε*_. To construct a travelling pulse solution on ℳ_*ε*_, we study the unstable and stable manifolds of 𝒞_0_ in their intersection with ℳ_*ε*_; depending on the sign of *c*, this effectively means we disregard the fast expansion (*c* > 0) or contraction (*c* < 0) in the (normal) *s*-direction. Defining the 3-dimensional manifolds

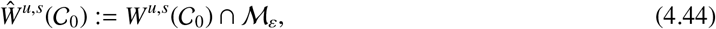

we want to determine whether *Ŵ ^u^*(𝒞_0_) and *Ŵ ^s^*(𝒞_0_) intersect transversally, in order to repeat the construction procedure outlined in subsection 4.1, see also [12, 13, 14, 48]. From the existence of a planar homoclinic orbit in the fast singular limit (4.40) (see subsection 4.1.1), it follows that the singular limits of *Ŵ ^u^*(𝒞_0_) and *Ŵ ^s^*(𝒞_0_) coincide, as in the stationary case. However, due to the fact that the full system (4.37) does *not* exhibit reversibility symmetry due to the presence of the *c*-dependent terms, the question whether *Ŵ ^u^*(𝒞_0_) and *Ŵ ^s^*(𝒞_0_) intersect transversally for nonzero *ε* cannot be answered based on symmetry arguments alone, as was the case in section 4.1.3, see also [13, 48]. Here, we need to perform a Melnikov calculation, along the lines of the analysis in [13, 48]. The Melnikov calculation is based on the observation that the singular limit system (4.40), in particular its planar reduction (4.15), is Hamiltonian. The associated conserved quantity *H*_*f*_ (4.18) is by definition zero along the singular planar homoclinic *w*_*h*_(*y*), as *w*_*h*_ lies on the level set *H*_*f*_ = 0; moreover, we see that *H*_*f*_ identically vanishes on 𝒞_0_. Hence, the total change of *H*_*f*_ accumulated along any orbit in *Ŵ ^u^*(𝒞_0_) ∩ *Ŵ ^s^*(𝒞_0_) (which is necessarily biasymptotic to 𝒞_0_), must vanish.

In terms of the original system variables (*u, p, v, q, s*), *H*_*f*_ takes the form

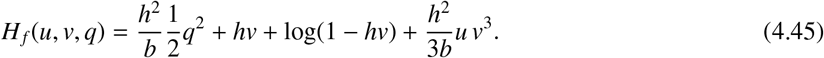

Hence, the change of *H*_*f*_ in *z* is given by

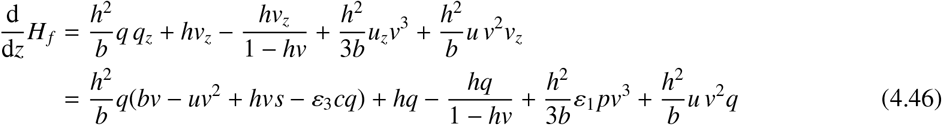

by (4.37). Using (4.37e), we can write

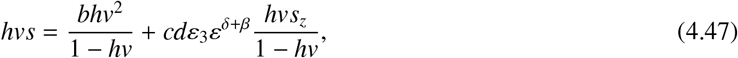

so that we obtain

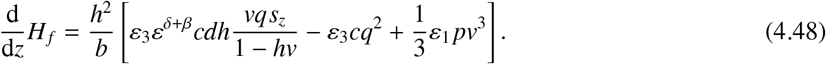

Following the analysis in [14, 48], we use the previously defined interval *I* _*f*_ (4.27) to express the total change of *H*_*f*_ accumulated along an orbit in *Ŵ ^u^*(𝒞_0_) ∩ *Ŵ ^s^*(𝒞_0_), as

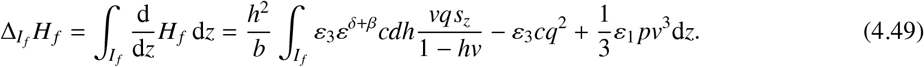

Now, we can use previous analysis on the reduced system (4.42) to obtain leading order expressions for (*v, q, s*), which can be used to obtain a leading order expression for Δ_*I*_*f H*_*f*_. Defining the positive integrals

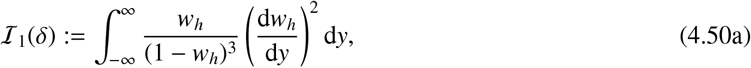

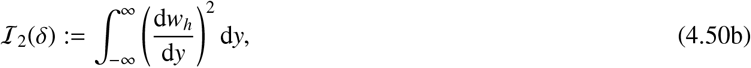

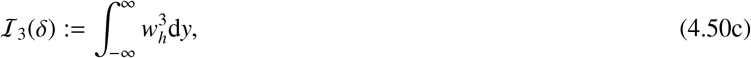

we can use (4.10), (4.16), (4.28) and (4.29) to obtain the condition

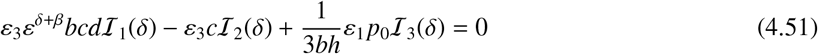

for Δ_*I*_*f H*_*f*_ to vanish to leading order. Note that the *h* → 0 (i.e. *δ* → 0) limit of (4.51) can be calculated using the asymptotic expansion (4.23)-(4.24), yielding

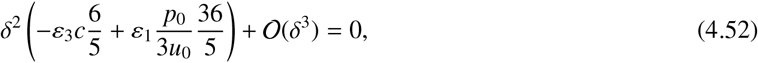

which is equivalent to [13, equation (3.14)] and [48, equation (2.24)].

As in the stationary case (cf. subsection 4.1.3), the second step in the construction of a pulse solution is to match the fast evolution through *Ŵ ^u^*(𝒞_0_) ∩ *Ŵ ^s^*(𝒞_0_) with the evolution on 𝒞_0_, such that the fast excursion normal to 𝒞_0_ can be concatenated with the slow flow on 𝒞_0_ along *E*^*u,s*^ (4.39), in order to construct an orbit that is homoclinic to the equilibrium point 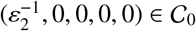. To that end, we need to calculate the change of the slow componentents (*u, p*) during the fast excursion. Since the dynamical equations for *u* and *p* in (4.37) differ from those in the stationary system (4.8) only by an asymptotically small term *ε*_3_*ε*^2^*cp*, and the leading order fast flow of (4.42) is the same as that of the stationary system (4.8), the leading order calculations (4.30) apply in the current setting as well. However, as the unstable and stable manifolds of the slow flow (4.39) differ from those in the stationary case (4.26), the resulting matching condition is different, and yields

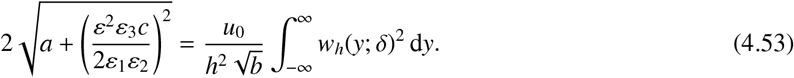

Note that at this point, the asymptotic magnitude of the fraction 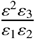 is still undetermined. However, as the right hand side of the matching condition (4.53) is 𝒪 (1) in *ε* and *ε*_1,2,3_, the left hand side must be as well, hence it follows that either 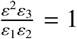 or 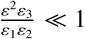.

Moreover, we can calculate the value of *p*_0_ in the existence condition (4.51) using (4.29), by writing

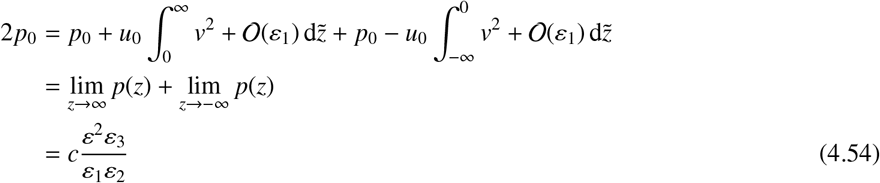

by (4.39); note that this is the same calculation as found in [48, equations (2.22-25)] and [13, equation (3.18)]. However, once we substitute this result into (4.51), we obtain the leading order existence condition

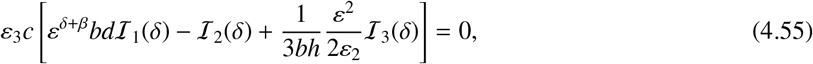

where we observe that *c* occurs as a (nonzero) prefactor; hence, the leading order value of *c* cannot be determined from (4.55). It follows that *c* must be determined from (4.53); to be able to do this, we see that 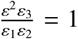, and thus *c* is the solution of

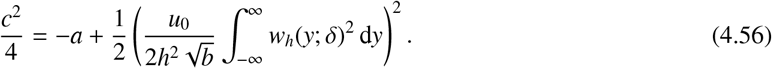

Moreover, to satisfy (4.55), we see that *ε*^*δ*+*β*^ or 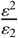 must be equal to one, since all integrals ℐ_1,2,3_ are positive and at least one between the first (ℐ_1_) and the third (ℐ_3_) term must balance ℐ_2_; from the same argument, it follows that neither term can be asymptotically large, hence *ε*^*δ*+*β*^ ≤ 1 and 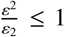. Thus, three distinct situations arise, where only the first (ℐ_1_) term, only the third (ℐ_3_) term, or both the first and the third term balance the second (ℐ_2_) term in the existence condition (4.55) In Figure 12, several parameter-dependent solution curves for these three different scaling choices of *ε*^*δ*+*β*^ and 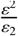 are shown.

**Figure 11:**
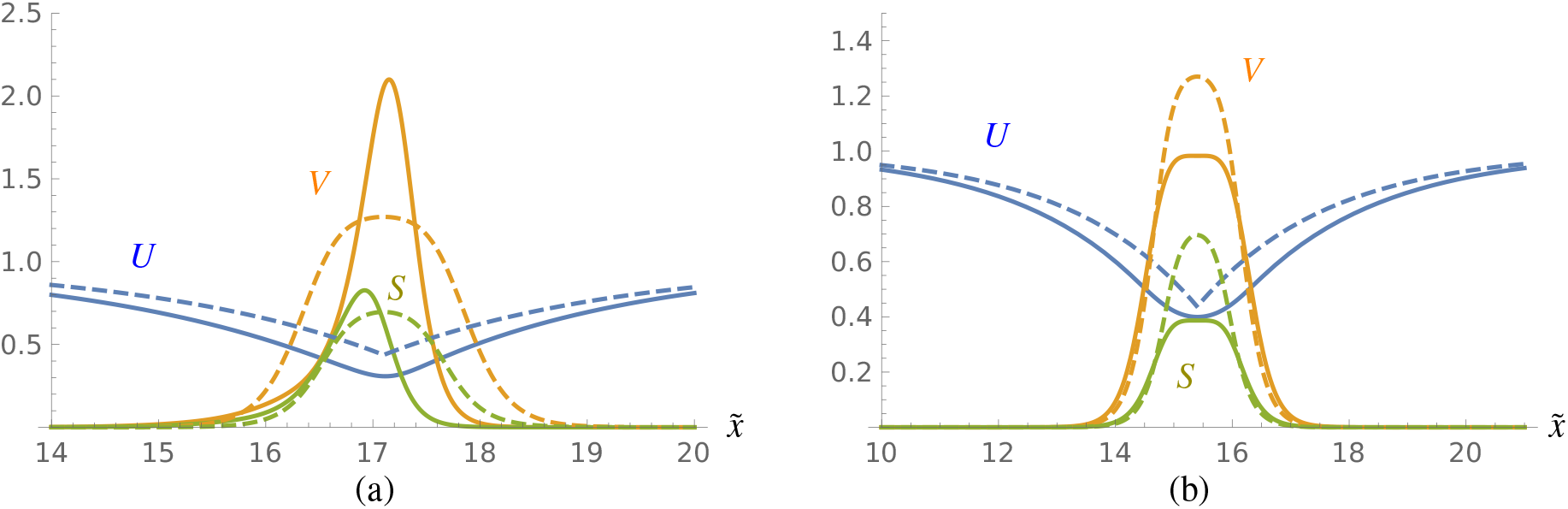
A comparison between direct numerical simulations of (4.1) (cf. Table 2) and the leading order analytical expression for a stationary pulse (see Theorem 4.3 and Figure 10 (b)), for parameter values 𝒜= ℬ= 0.2, ℋ= 0.5 and *ε* 𝒟= 0.1, with *U* in blue, *V* in orange and *S* in green. (a) The travelling pulse profile that is observed for = 4.5, solid; the leading order analytical stationary pulse solution, dashed. The differences between the pulse profiles suggest that the numerically observed travelling pulse is not a straightforward perturbation of the analytical stationary pulse. (b) The stationary pulse profile that is observed for 𝒟 = 0.4, solid; the leading order analytical stationary pulse solution, dashed. The difference between the pulse profiles are of 𝒪 (*ε*); the profiles are seen to match more closely when *ε* is decreased.

**Figure 12:**
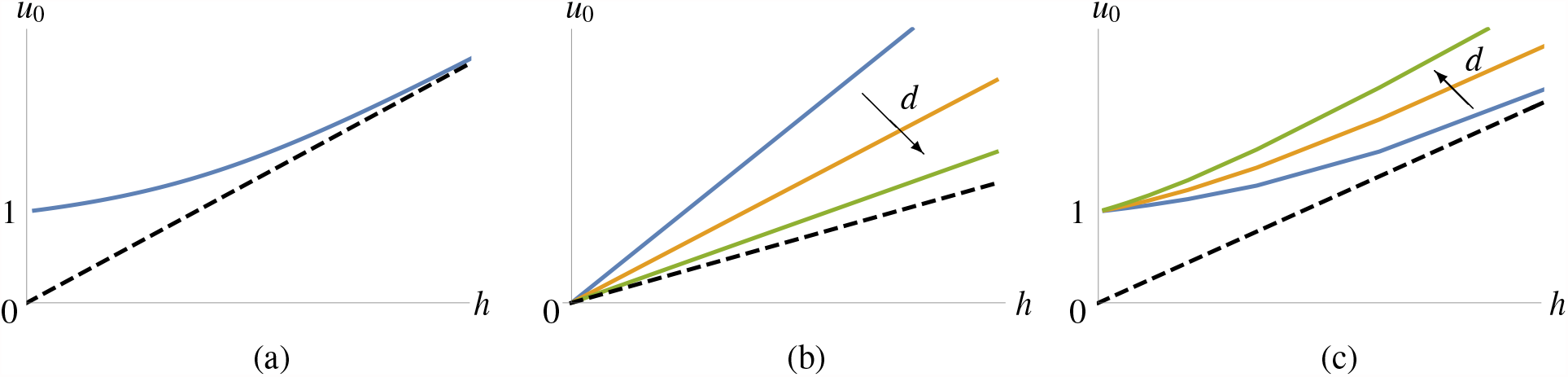
Solution curves of the existence condition (4.55), for different scaling choices of *ε*^*δ*+*β*^ and 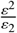, where *a* and *b* are fixed throughout. The set of admissible values, for which 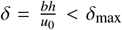, is bounded below by the dashed line 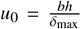. (a) Dominant balance between the second (ℐ_2_) and third (ℐ_3_) term, i.e. 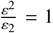 and *ε*^*δ*+*β*^ ≪ 1. Here, *u*_0_ is independent of *d*. (b) Dominant balance between the first (ℐ_1_) and the second (ℐ_2_) term, i.e. 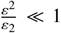 and *ε*^*δ*+*β*^ = 1. Here, *u*_0_ is plotted as a function of *h* for increasing values of *d*. (c) Equal balance between all three terms, i.e. both 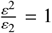 and *ε*^*δ*+*β*^ = 1. Again, *u*_0_ is plotted as a function of *h* for increasing values of *d*.

We summarise the findings of this subsection in the following theorem:

##### **Theorem 4.5**.

*Let ε be sufficiently small. Moreover, let* 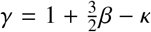, *and assume at least one of the following conditions holds:*

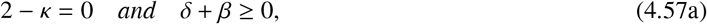

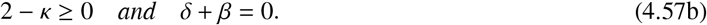

*Then, system* (4.37) *admits a homoclinic orbit to the equilibrium* 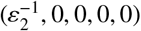, *provided equation* (4.56) *admits a real solution for c, where δ is chosen such that condition* (4.55) *is satisfied to leading order in ε*.

*The homoclinic orbit is, to leading order in ε, given by*

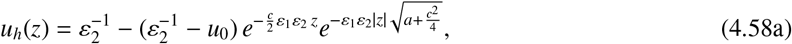

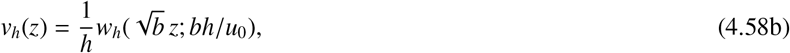

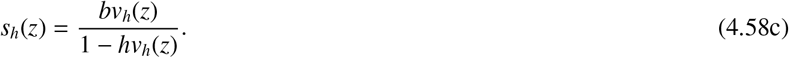

*Conversely, if either* (4.56) *has no solution for real c, or* (4.55) *cannot be satisfied, then no orbit homoclinic to* 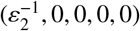 *exists*.

##### *Remark* 4.6

(*Non-existence of travelling pulses in Gray-Scott*). The existence result of Theorem 4.5 does not contradict the non-existence result [13, Theorem 5.1] for travelling pulses, as the scaling regimes (4.57a), (4.57b) were not considered in [13]; see also [12, Remark 4.1].

*Proof*. For sufficiently small *ε*_3_*ε*^*δ*+*β*^, Fenichel’s invariant manifold theorem [22, 30] provides the existence of ℳ_*ε*_, which is invariant under the flow of (4.37) and 𝒪 (*ε*_3_*ε*^*δ+β*^) close to ℳ_0_ (4.12). Specifically, there exists *ε*_0,*A*_ > 0 such that for all 0 < *ε* < *ε*_0,*A*_, the persistence of ℳ_*ε*_ as an invariant manifold is guaranteed. On this 4-dimensional invariant manifold ℳ_*ε*_, the transversal intersection of the restricted unstable and stable manifolds of _0_ (4.44), *Ŵ ^u^*(𝒞_0_) ∩*Ŵ ^s^*(𝒞_0_), follows from leading order calculations presented in subsection 4.2.2 combined with geometric arguments from the proof of [13, Theorem 4.1] and/or [48, Theorem 2.1]. The latter only needs to be augmented in the construction of the take-off and touchdown curves, where the intersection with the hyperplane {*q* = 0} now needs to be considered in its intersection with ℳ_*ε*_. Geometric perturbation theory now provides the existence of *ε*_0,*B*_ > 0 such that for all 0 < *ε* < *ε*_0,*B*_, a homoclinic orbit to 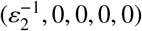 exists within ℳ_*ε*_; defining the global upper *ε*-bound 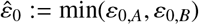 completes the proof.

It is useful to explore the ramifications of the scaling choices made in this subsection for the original system (4.1). As mentioned in Theorem 4.5, the condition 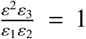 fixes 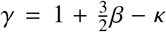, which, in combination with condition (4.6), implies *β* κ > 0.

The scaling choice (4.57a), which balances ℐ_2_ with ℐ_3_, implies *β* > 2. Taken together with κ = 2, we find for the scaling exponents of the original model parameters (4.5):

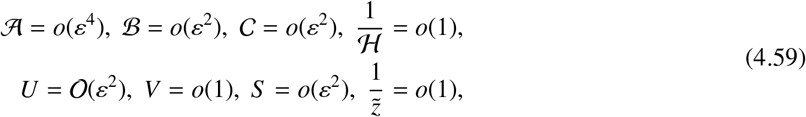

while the sign of *δ*, the scaling exponent of 𝒟, is not fixed. In particular, we see that 𝒜 scales with a relatively high power of *ε*; hence, a travelling pulse whose existence is established by Theorem 4.5 with scaling (4.57a), can only be observed for very small values of 𝒜. In addition, the *h* → 0 limit of (4.55) yields *u*_0_ = 1 and 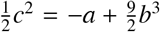 (see also Figure 12 (a)). That is, in the classical KGS limit, the amplitude of the pulse does, to leading order, *not* depend on the system parameters; this is in clear contrast to the stationary pulses studied in this paper (see Theorem 4.3) and to the travelling pulses constructed in [48].

On the other hand, the scaling choice (4.57a), which balances ℐ_2_ with ℐ_1_, only implies *β* > 0. Taken together with 0 < κ ≤ 2, we find for the scaling exponents of the original model parameters (4.5):

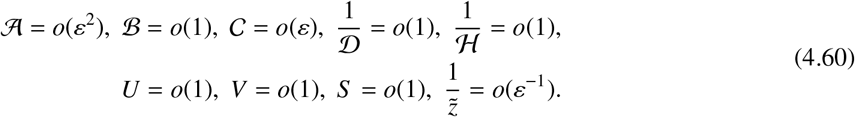

Compared with (4.59), the conditions on the scaling components are rather mild. Still, the condition *β* − κ > 0 implies that 𝒪 (𝒜/ ℬ) = *o*(*ε*^2^) – that is, 𝒜 and ℬ differ at least by two orders of magnitude in *ε*. Note that the *h* → 0 limit of (4.55) yields *u*_0_ = 0 (see also Figure 12 (b)), accentuating the fact that this solution branch does not exist in the classical KGS system.

#### 4.2.3. Constructing a travelling pulse solution, case II: (v, q, s)-dynamics on the same scale

We assume that the *s*-dynamics occur on the same scale as the (*v, q*)-dynamics, that is, *ε*_3_*ε*^*δ+β*^ = 1. Taking the limit *ε* → 0 in (4.37) now yields

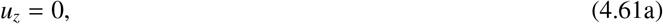

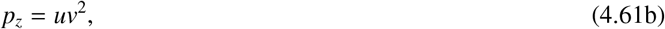

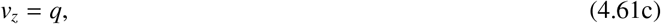

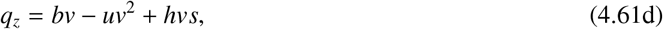

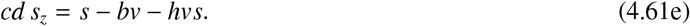

The construction method of the previous sections can be applied if and only if the reduced system (4.61) admits a homoclinic orbit. In contrast to the previous cases (cf. (4.15) and (4.40)), where the effective phase space was planar, the dynamics of (4.61) are effectively three-dimensional. This poses a considerate analytical challenge, as the dynamics of nonlinear three-dimensional dynamical systems can be highly complex. That being said, it seems sensible to start from a situation where the existence of a homoclinic orbit is known – that is, in the singular limit *cd* = 0. In this limit, the analysis of section 4.1.1 applies, and provides us with an homoclinic orbit (*v*_*h*_(*z*), *q*_*h*_(*z*), *s*_*h*_(*z*)). Now, we can use Melnikov theory [24, section 2.1] to determine whether the intersection of the stable and unstable manifolds of the origin (*v, q, s*) = (0, 0, 0) persists for *cd* > 0. To that end, we use the conserved quantity *H*_*f*_ in terms of the original variables (*v, q*) (4.46). However, we find that

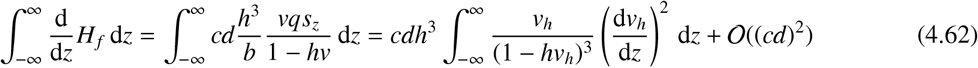

is always positive, since *v*_*h*_ > 0 by assumption, and 1 − *hv*_*h*_ > 0, see (4.11). This suffices to conclude that the intersection for *cd* = 0 does not persist for small positive values of *cd*.

While, in principle, the existence of homoclinic orbits in (4.61) for non-small values of *cd* is not ruled out by the above argument, their analytical inaccessibility makes the situation created by the scaling choice considered in this section, fall outside the scope of this paper. Hence, we disregard the (limiting) case *ε*_3_*ε*^*δ+β*^ = 1.

#### 4.2.4. Constructing a travelling pulse solution, case III: s slower than (v, q)

We assume that the *s*-dynamics are slower than the (*v, q*)-dynamics, that is, 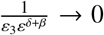 as *ε* → 0. Multiplying (4.37e) with 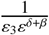 and subsequently taking the limit *ε* → 0 in (4.37) then yields

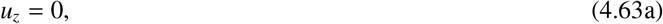

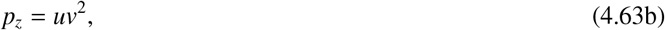

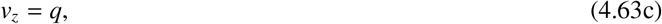

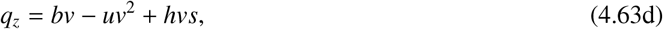

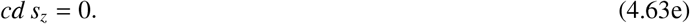

The existence of a homoclinic orbit in (4.63) now follows directly from the analysis in [13, 48], yielding

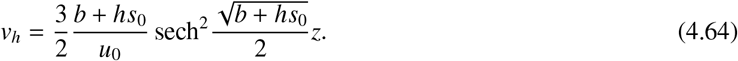

In principle, one could mimic the analytical steps taken in subsection 4.2.2, to obtain existence conditions along the lines of (4.55), which would depend on *s*_0_. However, for reasons that will become clear below, we will consider the dynamics of the slower variables (*u, p, s*) first.

The set of equilibria of the reduced system (4.63) is given by the 3-dimensional hyperplane

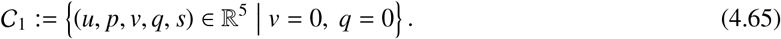

Like 𝒞_0_ (4.14), 𝒞_1_ is invariant under the flow of the full 5-dimensional system (4.37). The (*u, p*)-dynamics on 𝒞_1_ are the same as those on 𝒞_0_, and are therefore given by system (4.25). However, the *s*-dynamics on 𝒞_1_ are given by

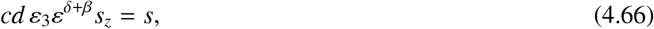

which are unbounded for z → ∞. Hence, we can conclude that in the scaling chosen in this subsection, any orbit homoclinic to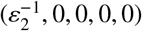 in (4.37), being necessarily bounded, must have a trivial *s*-component – that is, must be a solution to the classical KGS system.

## 5. Conclusion and outlook

The aim of this paper is to analytically investigate the influence of autotoxicity on vegetation patterns, through the analysis of travelling pulse solutions to the biomass-water-toxicity model (2.1). In particular, we want to analytically explain the occurence of travelling, asymmetric pulses, as observed in numerical simulations of the biomass-water-toxicity system, see e.g. Figure 3. The numerical evidence presented in section 3 suggests that the presence of toxicity is a prerequisite for the occurence of these travelling pulses in particular (compare Figures 2 and 3), and for previously unobserved dynamic spatio-temporal patterns in general (cf. Figure 1).

However, while the inclusion of an additional model component to the classical KGS model enriches the class of spatial patterns produced by this model with a number of provocative spatiotemporal patterning phenomena, it also significantly increases the mathematical complexity of the model, and subsequently of its analysis. Where the occurence of several types of patterns in the classical KGS model sensitively depends on the asymptotic scaling of the model components and parameters (see e.g. [12, Figure 2]), this is even more so the case in the extended model (2.4) considered in this paper. Therefore, the main emphasis of the analytical part of this paper (section 4) is on the balance of asymptotic scalings. The result of this analysis is that, within the scaling preparatory scaling choices made in AppendixA, stationary and travelling pulses can only be constructed in specific asymptotic scalings. This ‘positive’ outcome, as summarised in Theorems 4.3 and 4.5, is balanced by the ‘negative’ outcome of subsections 4.2.3 and 4.2.4.

Comparing the statement of Theorem 4.5 and its ramifications for the parameter scalings in the nondimensionalised model (2.4), it is clear that the travelling pulses constructed in subsection 4.2.2 are *not* the travelling pulses observed in numerical simulations, shown in Figure 3. In particular, the numerically observed pulses vanish when the parameters 𝒜 and ℬ are not approximately equal (see Table 2), whereas the travelling pulses from Theorem 4.5 only exist when 𝒜 and ℬ differ by two orders of magnitude in *ε* (cf. (4.59) and (4.60)). This discrepancy between analytical results and numerical observations could be explained in several ways.

- **Stability properties**. As mentioned in section 3, our numerical investigation indicates that the travelling pulse observed in the simulations for specific parameter ranges is stable in the PDE sense. Since the travelling pulses constructed analytically in section 4 are not recoverable in our simulations – i.e., even when the parameter constraints in (4.59), (4.60) are obeyed and/or we consider the leading order approximation of the pulse in (4.58) as initial condition the system converges to a uniform steady state and the patterned structure breaks down – we expect the analytically constructed pulses to be unstable in the PDE sense.
- **Boundary e**ff**ects**. The numerical simulations have been carried out on a bounded domain, whereas the analytical results assume an unbounded spatial domain. Hence, the existence of travelling pulses could be accredited to boundary effects – in particular, to self-interaction through the boundary. This would mean that a single travelling pulse on an unbounded domain is not the proper mathematical abstraction of the numerical observations as in Figure 3. Rather, one should consider a pair of interacting pulses, along the lines of [12]. However, our numerical investigations reveal that the observed travelling pulse solutions persist (both shape and speed) when doubling of the domain length, which is not in line with the hypothesis of self-interaction (see Remark 3.1).
- **Transient behaviour**. The numerical simulations have been carried out for a finite time, whereas the existence analysis is time-independent. Hence, the observed travelling pulses could be the manifestation of a slow transient process from initial state to bare soil, where these travelling pulses only exist for a finite time. This would mean that a time-independent travelling pulse, that is, a stationary pulse in a co-moving frame with fixed speed, is not the proper mathematical abstraction of the numerical observations as in Figure 3. However, our numerical investigations reveal that the observed travelling pulse solutions persist when doubling the simulation time (see Remark 3.2). This does not rule out the hypothesis of metastability, but it does severely limit the evolution speed of the wave profile. Transitional pattern formation phenomena have be studied analytically, see e.g. [3]; one could adopt the approach outlined in [3] to study (2.4). However, the absence of integrable structure might prohibit this approach.
- **Scaling assumptions**. As outlined in the introduction of section 4, the applicability of the geometric singular perturbation theory approach to pulse patterns, as used in [12, 13, 14, 48], is closely tied to the asymptotic scaling of the underlying model (2.4). The arguments leading to the preparatory scaling choices (4.5) are specified in AppendixA. However, it is important to note that not every scaling choice made in AppendixA is *necessary* for the application of the geometric construction techniques. The two scaling choices pertaining to the dynamics of the toxicity component *s* ((A.17) and (A.20)) are convenient, rather than necessary. In particular, the assumption underlying the last scaling choice (A.20) – namely, that the stationary pulse constructed in section 4.1 should be the ‘*c* = 0’ member of a family of travelling (‘*c* * 0’) pulses – could prove to be too restrictive. After all, families of travelling waves that do not include a neighbourhood of *c* = 0 regularly occur [8, 51, 53]. In addition, the ‘proper’ scaling needed to understand the travelling pulse shown in Figure 5 need not be uniform: abandoning scalings (A.17) and (A.20) might lead to the situation where the *nonlinear* interaction terms govern the singular behaviour of the pulse solution. In such a case, one would need to use geometric blowup techniques to construct a singular concatenated homoclinic orbit, along the lines of [25].

The analysis of travelling pulses incorporating this generalised scaling is ongoing work.

It is important to note that, while the analytical approach advocated in this paper applies to the *existence* of stationary and travelling pulse solutions, the question of pulse *stability* is still very much open. We plan to apply the techniques developed for the stability analysis of pulses in two-component reaction-diffusion systems, as presented in [14], to the three-component reaction-diffusion-ODE system (2.4). The method presented in [14], which is based on an Evans function approach, does not in principle depend on the number of model components, and can be applied to pattern solutions in *n*-dimensional reaction-diffusion systems [47]. Moreover, this approach is amenable to be extended to systems of mixed (reaction-diffusion-ODE) type. The stability of pulse solutions to the biomass-water-toxicity model (2.1) will be investigated in a separate work.

Moreover, numerical simulations in [35] on two-dimensional domains reveal the presence of both crescent (travelling) moon spots and double-scale patterns for some parameter regimes. In these double-scale pattens, the pulses/fronts travel at a micro-scale level within a pattern, which appears stable on a macroscopic scale. Such multiple-scale behaviour has been connected with pattern robustness in a different, but related, ecological context [33]. Future challenges hence involve the analytical investigation of such structures – in the first case (travelling crescent moons), an approach along the lines of [21] would be a prime candidate.

The biomass-water-toxicity model by Marasco et al. (2.1) has proven to be a rich and inspiring source of previously unobserved patterning phenomena. It is important to emphasise that autotoxicity can be used to explain and recreate experimentally observed dynamical patterns, without having to assume a specific domain topography, in contrast to previous work [1, 2, 52]. In this respect, the model is interesting both from an ecological perspective, and from the more general viewpoint of mathematical modelling. Systems of reaction-diffusion-ODE type have been the subject of recent investigations; in particular, the shape and stability properties of patterns have proven to be significantly different from their ‘classical reaction-diffusion’ counterparts, sometimes leading to counterintuitive results [36, 37]. We hope that the work presented in this paper, though exploratory, will lead to a deeper understanding and a broader appreciation of systems of this type, of which the biomass-water-toxicity model (2.1) is an intriguing example.

## Acknowledgements

AI acknowledges partial support from the New Frontier’s grant NST-0001 of the Austrian Academy of Sciences ÖAW and an FWF Hertha Firnberg Research Fellowship (T 1199-N). FV was supported by a Humboldt Research Fellowship.

## AppendixA Preparatory asymptotic scaling of system (4.4)

Our goal is to scale system (4.4) in such a way, that we can apply the approach taken in e.g. [13, 14, 48] to construct a travelling pulse solution. For similar asymptotic scaling analyses (with the same goal), see [13, Appendix] and [48, section 2.1].

A priori, every parameter, component, and variable in system (4.4) admits an asymptotic scaling in the small parameter ε. Hence, we scale

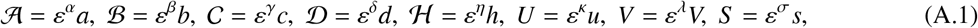

and introduce a rescaled coordinate 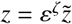 to obtain

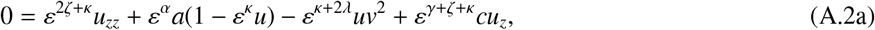

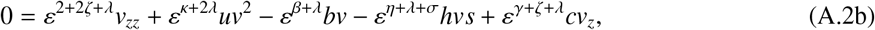

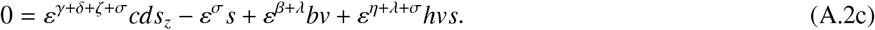

We now choose *z* to be the variable on which the (fast) *v*-dynamics take place. Moreover, we assume that the equilibrium (*u, v, s*) = (*ε*^*−κ*^, 0, 0) is hyperbolic in *v*. This fixes the asymptotic scaling of the coordinate *z* as

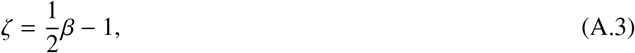

which, dividing the second equation in (A.2) by *ε*^*β*+λ^, yields

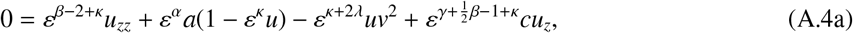

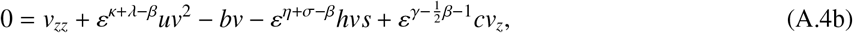

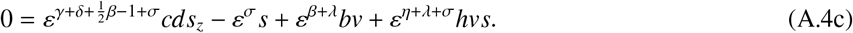

In the geometric construction as carried out in [13, 14, 48], a pivotal element is the existence of a homoclinic orbit (spike) in *v* for a fixed value of *u*. In particular, this means that the initial exponential growth close to *v* = 0 due to the term −*bv* in (A.4b) must be balanced by a positive, nonlinear term. We fix the asymptotic scaling of *v* such, that such an homoclinic spike solution has 𝒪 (1) amplitude in *v*, which implies

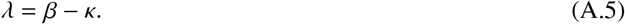

With this scaling, dividing the third equation in (A.4) by *ε*^σ^, we obtain

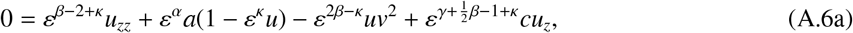

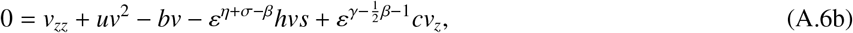

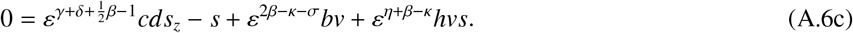

Furthermore, based on the numerical observations from section 3, we conjecture that a homoclinic spike solution in *v* is, to leading order, symmetric in *z*. This implies that the advection term *cv*_*z*_ in the *v*-equation is perturbative, hence

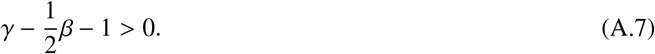

Next, we consider the behaviour of *u*. For the type of pulse solutions we consider in this paper, there is a clear spatial scale separation between the *u*-component and the *v*-component, see Figure 5. In particular, *u* is *slow* in *z* in comparison to *v*. Rewriting (A.6a) as

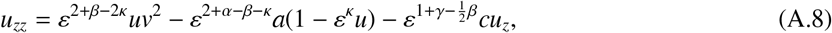

this implies that

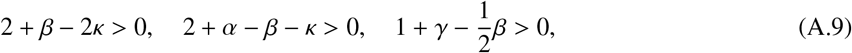

where the third inequality already follows from (A.7). Moreover, both from the simulation results presented section 3 and the interpretation of the model (2.4) in light of its dimensional version (2.1), it is clear that the *U*-variable takes values *in between* the homogeneous dry (*U* = 0) and the homogeneous wet (*U* = 1) states. As *U* = *ε*^κ^*u*, this implies

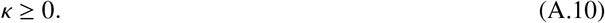

For presentation purposes, we henceforth exclude the case *κ* = 0, to avoid having to treat several scaling subcases independently. However, the analysis in section 4 covers the case *κ* = 0 as well, as it is nowhere necessary to assume that *κ* > 0; see also [48, Remarks 1 and 2].

The coupling of *v* into the *u*-equation plays a central role in the geometric construction outlined in section 4 of this paper, and in the main references [12, 13, 14, 48]. In particular, the question is whether the stable and unstable manifolds of the invariant manifold 𝒞 _0_ (4.14) intersect transversally, thereby guaranteeing the existence of a (travelling) pulse solution to (2.4). A necessary condition for this intersection to exist, is that the *uv*^2^-term in (A.8) is leading order – that is,

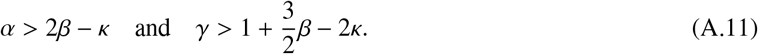

Moreover, this term should match, in a specific way, the dynamics of *u* on 𝒞 _0_ – that is, the dynamics of *u* when *v* = 0. From (A.8), we see that these dynamics are linear, with *u* = *ε*^−κ^ being the only equilibrium. This equilibrium is a saddle, with stable resp. unstable eigenvalues

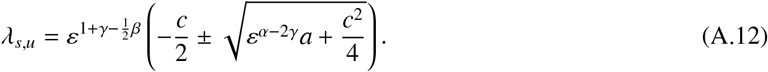

We now assume that there exists an asymptotic scaling such that both eigenvalues (A.12) are nonzero and of the same order, which implies

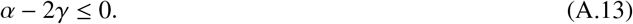

For the same eigenvalues, we consider the associated linear stable and unstable subspaces *E*^*s*^ and *E*^*u*^ in (*u, u*_*z*_)-phase space. The geometric matching condition now stipulates that the *vertical distance* between these two subspaces, which is given by

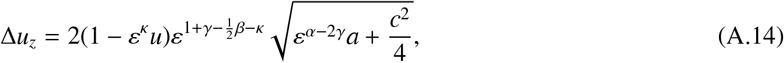

should be of the same asymptotic order as the leading order term in (A.8), that is, *ε*^2+*β*−2κ^*uv*^2^. We use this condition to fix the scaling of 𝒜, yielding

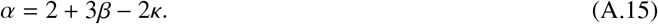

This scaling choice further specifies (A.6) as

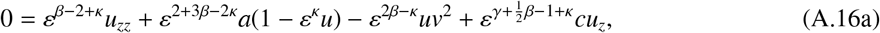

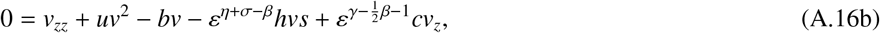

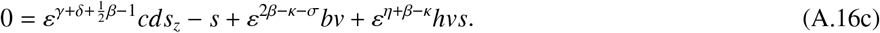

Next, we consider the aim of this paper – that is, to investigate the influence of toxicity on the dynamics of vegetation patterns. The coupling between the toxicity component *s* and the biomass component *v* is mediated by the term *hvs*. We fix the scaling of the parameter ℋ such, that the toxicity-coupling term has 𝒪 (1) influence on the biomass dynamics, which implies

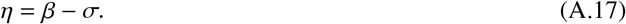

Applying this scaling choice to system (A.16), we obtain

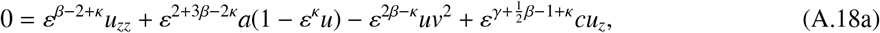

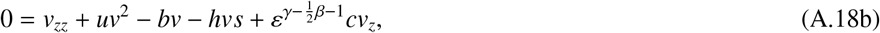

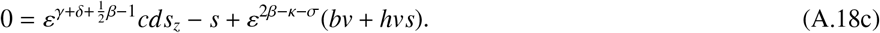

As a last scaling choice, we consider the question of existence of *stationary* pulse solutions – that is, when 𝒞 = 0 in (4.3), and as a consequence, *c* = 0 in (A.18). From the resulting (algebraic) *s*-equation

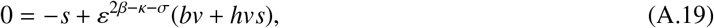

it follows that for a stationary pulse to have a nontrivial *s*-component, the terms in (A.19) all need to have the same asymptotic scaling. When one considers a stationary pulse solution to be a particular travelling pulse solution−namely, one with zero speed−it can be argued that both solution types have to exist within the same asymptotic scaling. Hence, we decide to fix the asymptotic amplitude of *s* as

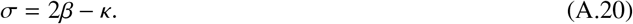

With this scaling, dividing the first equation of (A.18) by *ε*^*β*−2+κ^, we obtain

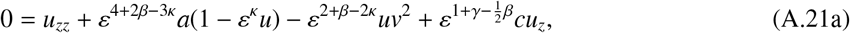

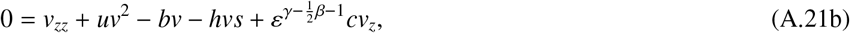

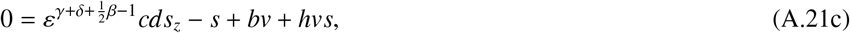

which is equivalent to the 5-dimensional system (4.7).

The first two equations of system (A.21) for *h* = 0 are equivalent to [13, system (A.3)]. Note that in [13], the additional scaling choice *A* = *ε*^2^*a* is made – that is, one scaling exponent is specifically chosen to be equal to two. In the context of the scaling presented in this appendix, this would be equivalent to the choice 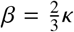. Note that this choice specifically precludes the travelling pulse solutions described in Theorem 4.5, see the discussion directly below this Theorem.

In addition, the first two equations of system (A.21) for *h* = 0 are equivalent to system [48, system (2.5)]. Note that, in [48], an additional scaling choice is made for the asymptotic scaling of the wave speed. In the context of the scaling presented in this appendix, this would be equivalent to the choice 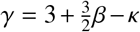, which implies *ε*_3_ = *ε*_1_, see (4.9), (4.36). This choice for the asymptotic scaling of the wave speed is different from the result 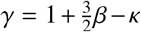 presented in Theorem 4.5; the two choices coincide only if κ = 2, which is a subcase of Theorem 4.5.

